# Parameter estimation and identifiability in a neural population model for electro-cortical activity

**DOI:** 10.1101/492504

**Authors:** Agus Hartoyo, Peter J. Cadusch, David T. J. Liley, Damien G. Hicks

## Abstract

Electroencephalography (EEG) provides a non-invasive measure of brain electrical activity. Neural population models, where large numbers of interacting neurons are considered collectively as a macroscopic system, have long been used to understand features in EEG signals. By tuning dozens of input parameters describing the excitatory and inhibitory neuron populations, these models can reproduce prominent features of the EEG such as the alpha-rhythm. However, the inverse problem, of directly estimating the parameters from fits to EEG data, remains unsolved. Solving this multi-parameter non-linear fitting problem will potentially provide a real-time method for characterizing average neuronal properties in human subjects. Here we perform unbiased fits of a 22-parameter neural population model to EEG data from 82 individuals, using both particle swarm optimization and Markov chain Monte Carlo sampling. We estimate how much is learned about individual parameters by computing Kullback-Leibler divergences between posterior and prior distributions for each parameter. Results indicate that only a single parameter, that determining the dynamics of inhibition, is directly identifiable, while other parameters have large, though correlated, uncertainties. We show that the eigenvalues of the Fisher information matrix are roughly uniformly spaced over a log scale, indicating that the model is sloppy, like many of the regulatory network models in systems biology. These eigenvalues indicate that the system can be modeled with a low effective dimensionality, with inhibition being prominent in driving system behavior.

**Author summary:** Electroencephalography (EEG), where electrodes are used to measure electric potential on the outside of the scalp, provides a simple, non-invasive way to study brain activity. Physiological interpretation of features in EEG signals has often involved use of collective models of neural populations. These neural population models have dozens of input parameters to describe the properties of inhibitory and excitatory neurons. Being able to estimate these parameters by direct fits to EEG data holds the promise of providing a real-time non-invasive method of inferring neuronal properties in different individuals. However, it has long been impossible to fit these nonlinear, multi-parameter models effectively. Here we describe fits of a 22-parameter neural population model to EEG spectra from 82 different subjects, all exhibiting alpha-oscillations. We show how only one parameter, that describing inhibitory dynamics, is constrained by the data, although all parameters are correlated. These results indicate that inhibition plays a central role in the generation and modulation of the alpha-rhythm in humans.

## Introduction

The classical alpha rhythm is one of the most remarkable features observed in electroencephalogram (EEG) recordings from humans [1, 2]. First discovered by Berger in the 1920s [3, 4] these waxing and waning oscillations of 8 - 13 Hz, that are prominent during eyes-closed, are a defining feature of the resting EEG and have played a central role in phenomenological descriptions of brain electromagnetic activity during cognition and in behaviour [5]. Despite being discovered almost a century ago, the alpha rhythm remains poorly understood, both in terms of its underlying physiological and dynamical mechanisms as well as its relevance to brain information processing and function. The received view proposes a central role for the thalamus [5] with early theories suggesting that alpha oscillatory activity intrinsic to thalamus ‘drives’ or ‘paces’ overlying cortical tissue [6]. This conception has been modified to suggest that it is feedback reverberant activity between thalamus and cortex which underpins the genesis of alpha band cortical activity [5, 7]. A contrasting hypothesis is that such oscillatory activity arises intrinsically in cortex, emerging purely from the recurrent activity between cortical populations of excitatory and inhibitory neurons. These different hypotheses have motivated a variety of theories for describing the alpha-rhythm [8–12].

Theories of alpha-rhythmogenesis can be divided into two major frameworks: those that take a nominal microscopic perspective by modeling the behaviour of large numbers of synaptically connected biophysically realistic individual neurons [8, 9] and those that take a macroscopic, or mean-field, stance by considering the activity of interacting *populations* of neurons [13–15]. While the microscopic approach is more fundamental, macroscopic models are better matched to the spatial scale at which the bulk electrophysiological measurement, the EEG, occurs.

Despite their reduced complexity, it is still extremely difficult to estimate the input parameters of neural population models by direct fits to EEG data. Up until now, the use of such models to explain alpha-rhythmogenesis has largely been limited to calculations of the *forward* problem: manually selecting input parameters such that the model generates alpha oscillations. It is vastly more difficult to solve the *inverse* problem, where a full set of parameters and their uncertainties are estimated directly from fits to EEG data. Yet solving this inverse problem is crucial if we are to ever regard the inferred parameter values as physiologically meaningful. As we will show, the fundamental challenge in fitting a neural population model is that many different combinations of input parameters can give the same EEG signal. Understanding the nature of such parameter unidentifiability (discussed next) in a neural population model is the major contribution of this paper.

### Parameter identifiability and sloppy models

Neural population models are high-order, multi-parameter, dynamical systems. It has long been known that, even in simple dynamical systems, there can exist very different parameter combinations which generate similar predictions [16–27]. This many-to-one mapping between parameter inputs and model observables is referred to as structural unidentifiability, if the predictions are exactly identical, or practical identifiability, if the predictions are nearly identical. Any fitting of an unidentifiable model to data results in large, correlated parameter uncertainties. Many developments in the study of identifiability in differential equation models have been motivated by problems in systems biology involving biomolecular regulatory networks [28–35].

Model unidentifiability is closely related to, though distinct from, model sloppiness. A model is referred to as sloppy if the sensitivity of its predictions for different parameters covers a broad range [36–41]. These sensitivities, quantified by the eigenvalues of the Fisher information matrix, are roughly uniformly spaced over a log scale. This characteristic has been discovered in a variety of nonlinear models and arises from the geometry of nonlinear models projected into data space [37, 38]. Parameters that sensitively affect model predictions are termed ‘stiff’ while those that can be changed with little effect on predictions are termed ‘sloppy’. While sloppy parameters are often unidentifiable as well, the terms are not synonymous [42, 43]. Like unidentifiability, sloppiness has been found to be prevalent in models of biomolecular networks [44–47].

Unidentifiability and sloppiness are pervasive in nonlinear fitting problems, the simplest examples of which are fits to polynomials or to multiple exponentials [36, 38]. Since parameter estimation in differential equation models always involves nonlinear fits (to exponential impulse responses in the time domain or rational transfer functions in the spectral domain, for example), unidentifiability and sloppiness should always be a concern in dynamical systems. This is true even for linear, time-invariant systems [19, 22, 26]. Of course, explicitly nonlinear functions of parameters certainly exacerbate the problem - a nonlinear function at saturation will give the same response for a range of different parameter inputs, for example. Unidentifiabilities also arise when a model supports phenomena at significantly different timescales. For instance, if only dynamics on a slower timescale can be observed, parameters which determine behavior at the faster timescale would not be constrained by data [48].

Unlike in systems biology modeling, in neurophysical modeling there has been little recognition of the problem of unidentifiability, beyond select examples in neural code models [49], a thalamo-cortical neural population model [50], and dynamic causal models [51]. This has been cited in [52] as an example of how approaches used in systems biology can help address problems in computational neuroscience [53]. However, despite this lack of formal discussion, implicit recognition of unidentifiability in computational neuroscience has been widespread, with several studies, including those for models of single neurons [54–58], occulomotor integration [59], and neural populations [60–62], detecting the large, correlated parameter uncertainties that are the hallmark of unidentifiability and sloppiness.

### Outline

In this paper we examine identifiability and sloppiness in a well-known neurophysical model [11, 61, 63, 64] with 22 unknown parameters. We concentrate on fitting EEG data exhibiting alpha-oscillations in resting state human subjects in an attempt to understand the mechanistic origin of this prominent, yet still poorly understood, phenomenon. We fit the model to the EEG spectrum from each of 82 subjects using both a particle swarm optimization and Markov chain Monte Carlo method. When viewed across all subjects, only 1 of the original parameters, the decay rate of inhibitory synaptic activity, emerges as being identifiable. This indicates that inhibition is essential for explaining alpha-rhythmogenesis. Examination of the Fisher information matrix shows that there are ~5 parameter *combinations* that are identifiable, a considerable reduction from the original 22. This indicates that, although most parameters are unidentifiable, their values cannot in general be selected arbitrarily to fit the data.

## Neural population model

The neural population model used in this paper is well established and has been described previously [11, 65]. Semi-analytical and numerical solutions of these equations have revealed a rich repertoire of physiologically plausible activity including noise driven, limit cycle and chaotic oscillations at the frequency of the mammalian alpha rhythm [11, 61, 66, 67]. Here we use the spatially homogeneous version given by the following coupled set of first and second order ordinary differential equations:

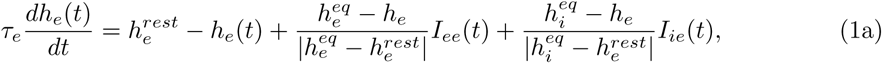

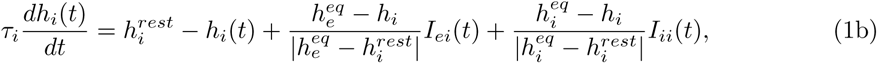

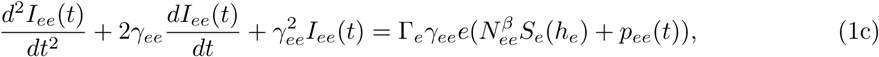

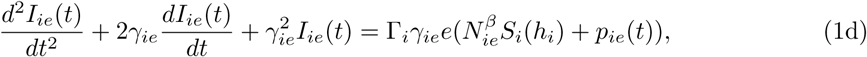

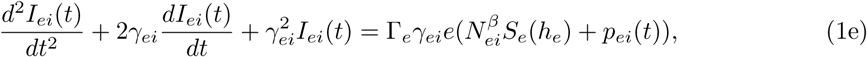

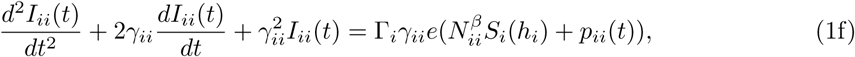

where

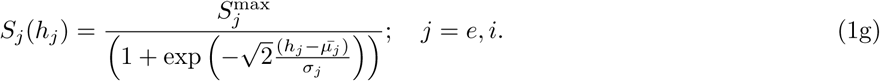

These equations describe the interactions between inhibitory and excitatory neuronal populations in a macrocolumn.

Table 1 lists the parameters along with their physiological ranges as assumed by Bojak and Liley [61]. The temporal dynamics of mean soma membrane potentials for the excitatory (*h_e_*(*t*)) and inhibitory (*h_i_*(*t*)) populations are described in Eq (1a) and (1b). The temporal dynamics of the synaptic activity, *I_ee_*(*t*), *I_ie_*(*t*), *I_ei_*(*t*), and *I_ii_(t)*, are given by Eqs (1c) - (1f). The relationship between the mean population firing rate, *S_j_*, and the mean soma membrane potentials of the respective population is given in Eq (1*g*). It has been shown that the local field potential measured in the EEG is linearly proportional to the mean soma membrane potentials of the excitatory populations, *h_e_*(*t*) [68, 69].

## Spectral analysis of EEG data

In this study we fit the above model to EEG recordings from 82 different individuals. This data is a subset of a larger dataset which, in its full version, consists of 14 experimental tasks performed by each of the 109 subjects with recordings on 64 electrodes according to the International 10-10 System. The full set, collected and contributed by Schalk et al [70] using the BCI2000 instrumentation system, is available for public access in PhysioNet [71] (https://www.physionet.org/pn4/eegmmidb/).

For the purpose of studying alpha-rhythm, we restricted our analysis to signals from the Oz electrode, selecting data from individuals whose EEG spectrum exhibited clear alpha peaks during the associated eyes-closed task. Welch’s method of averaging the spectra derived from multiple overlapping time segments [72] was used to estimate the single spectrum for a particular individual. This approach improves the precision of the power spectral density estimate by sacrificing some spectral resolution. A one-minute EEG signal associated with a particular individual, sampled at 160 Hz, was divided into segments using a 4-second Hamming window with an overlap of 50%.

Since the computational demands of fitting our model directly to EEG time series data are prohibitive, we fit the EEG spectrum instead. This approach is in accordance with earlier fits of neural population models [50, 61, 62], which involved fewer unknown parameters than we have here, and generally only fit a single EEG spectrum. We are thus assuming stationarity of the system over the one-minute EEG signal, where stationarity here means that it is the parameters that are constant; the states are allowed to vary about a stable fixed point. We furthermore assume that deviations of the state away from the fixed point are small enough to allow linearization of the model. Though parameters are assumed to be constant within a given EEG recording they can of course vary between different recordings (and thus individuals). Because of well-known nonlinearities and nonstationarities in EEG recordings, our linearized model was used in inference procedures only for frequencies between 2 Hz and 20 Hz [73].

To demonstrate the accuracy of our inference methods, we also show an example of parameter estimation from a simulated spectrum where the underlying parameter set is known (and referred to as the ground truth). In order to choose a plausible parameter set for this test, we use the maximum likelihood estimate found for Subject 77 (the estimate found from any other subject would also have been suitable). The simulated spectrum was then calculated by sampling each frequency channel from the gamma-distributed model prediction.

## Model fitting and analysis

To examine the identifiability and sloppiness of the neural population model, we fit to an EEG spectrum and estimate the posterior distribution over the 22 unknown parameters. We then characterize the properties of this distribution to diagnose the signatures of unidentifiability and sloppiness. To ensure that our results are not specific to a particular fitting algorithm, we use two independent methods: particle swarm optimization (PSO) and Markov chain Monte Carlo (MCMC). To ensure that our results are not specific to a given individual, we estimate the 22 posterior distributions, using both methods, for each of the 82 different EEG spectra.

A full description of the methods for fitting the data and analyzing the results is given in Section “Methods of analysis” where we first describe the procedure for calculating the predicted model spectrum, along with the likelihood function for the spectral estimate. We then outline the two fitting schemes, focusing on how we use them to sample from the 22-dimensional posterior distribution. Finally, we describe use of the Kullback-Leibler divergence (KLD) to summarize how much we learned about individual parameters, and the Fisher information matrix (FIM), to assess the sloppiness and identifiability of the model.

## Results

Fig 1 and Fig 2 illustrate best fits using PSO and MCMC methods. Although the fits are generally similar, subtle differences between the two methods can be observed. For example, in subject 72 the the ML fit performs better on regions with lower power but less well in regions with higher power. This difference is expected: while LS are computed over unweighted power spectra, ML favors frequencies with lower variances which typically are those with lower power spectra.

**Fig 1.**
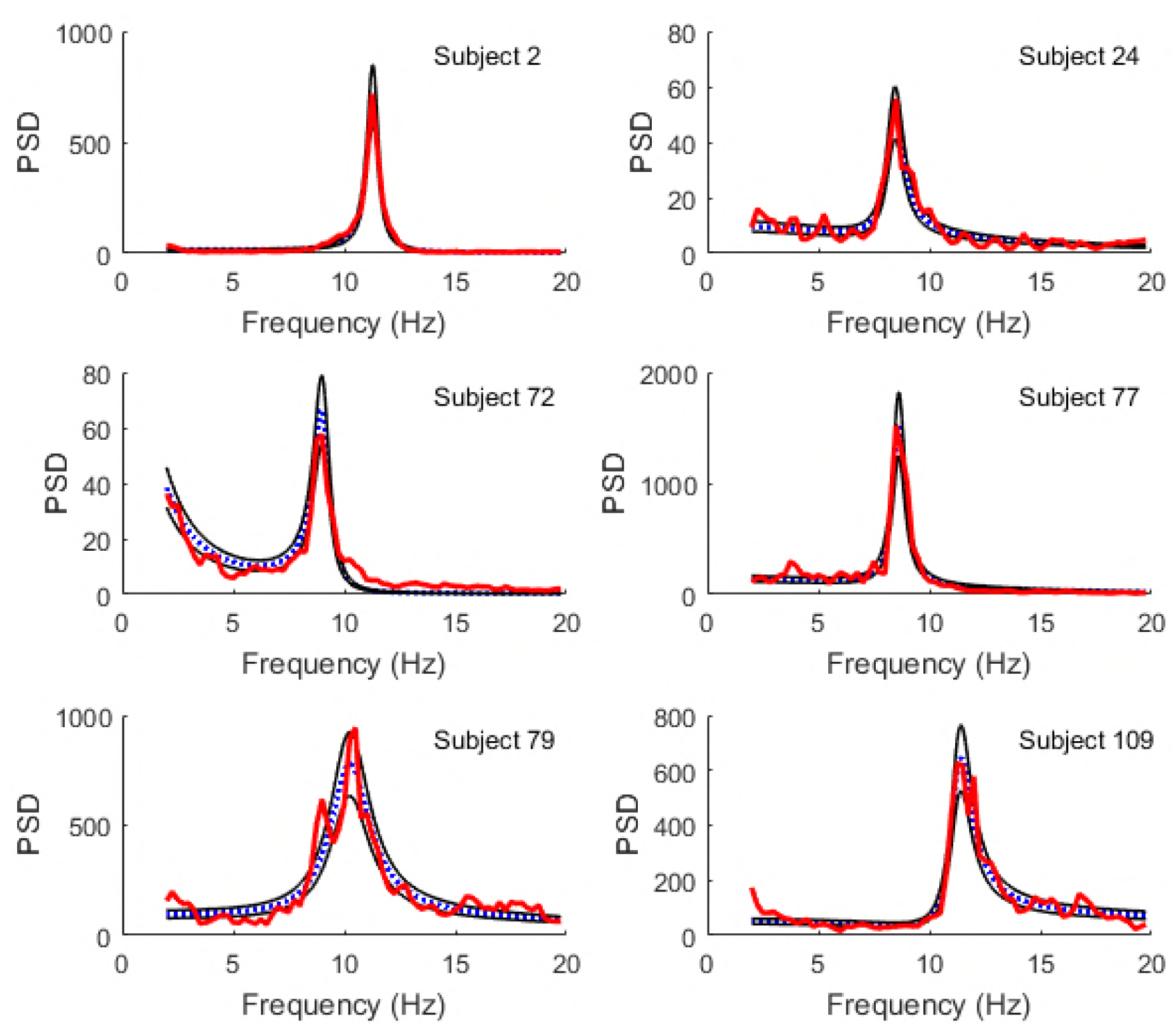
Best fits using least squares. Comparison of model spectra (blue dotted line) fit to experimental spectra (red thick line) by least squares (LS) minimization using particle swarm optimization, for a select set of subjects. Also shown are the 16% and 84% quantiles based on the gamma distribution for the fitted spectra (thin black lines). The subjects have been selected to show the range of spectra included in the full data set.

**Fig 2.**
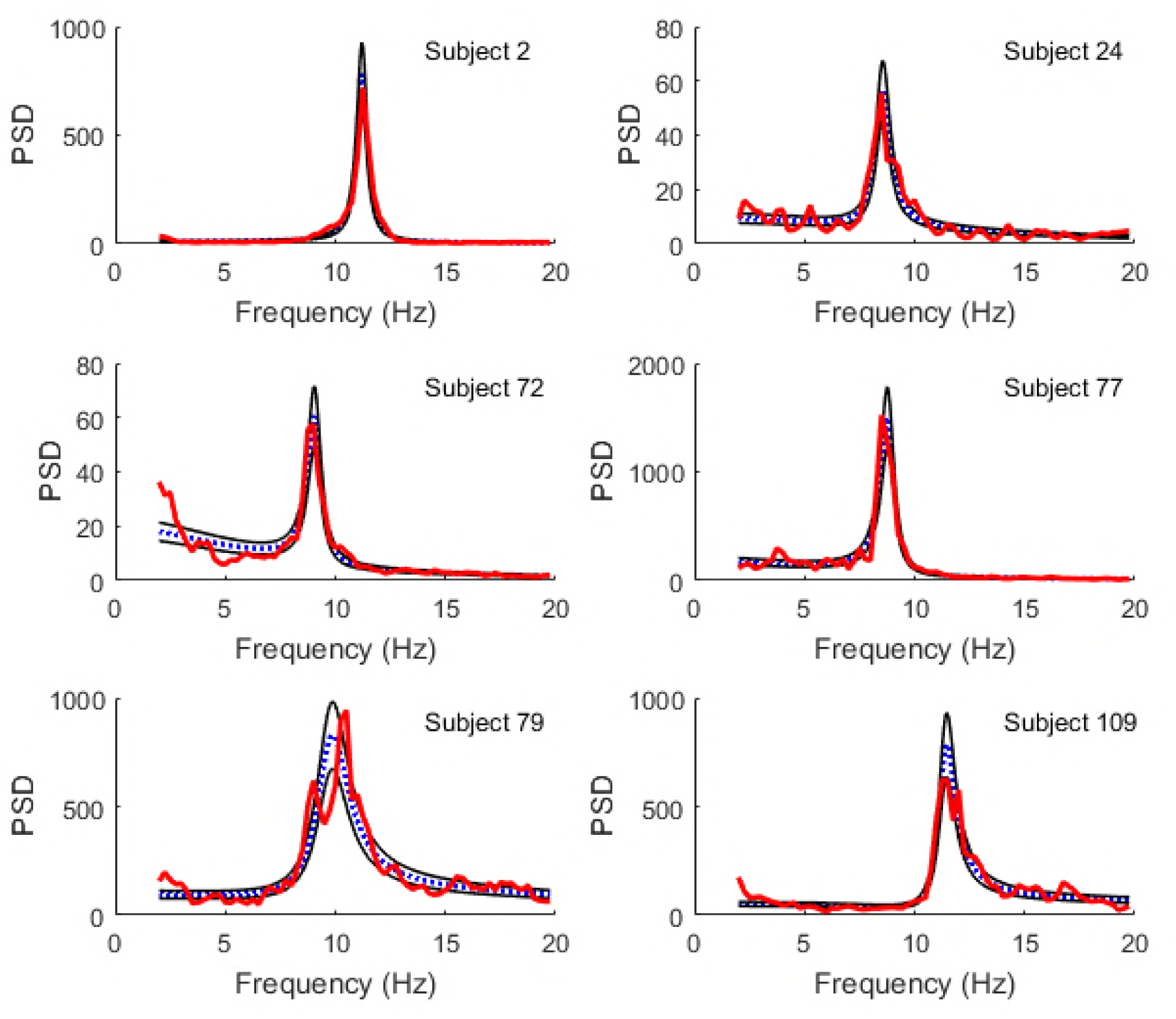
Best fits using maximum likelihood. Comparison of model spectra (blue dotted line) fit to experimental spectra (red thick line) by maximum likelihood (ML) estimation using MCMC. Also shown are the 16% and 84% quantiles (thin black lines). The subjects are the same as in Fig 1. It should be noted that LS and ML fits are expected to differ in this case, since the LS fits are more sensitive to deviations in the unweighted spectral power (typically in regions with larger power) whereas the ML fits are more sensitive to deviations in regions where the variance of spectral power is small (typically in regions of lower spectral power).

The posterior marginal distributions for each parameter, from a few subjects, are shown in blue in Fig 3 (from PSO sampling) and Fig 4 (from MCMC sampling). These are compared to the uniform prior distributions (green). The top row, which corresponds to analysis of the simulated spectrum, also shows the ground truth value (red). Each parameter is plotted in normalized coordinates, where -1 corresponds to the lower limit of the plausible parameter interval and +1 corresponds to the upper limit (see Table 1).

**Fig 3.**
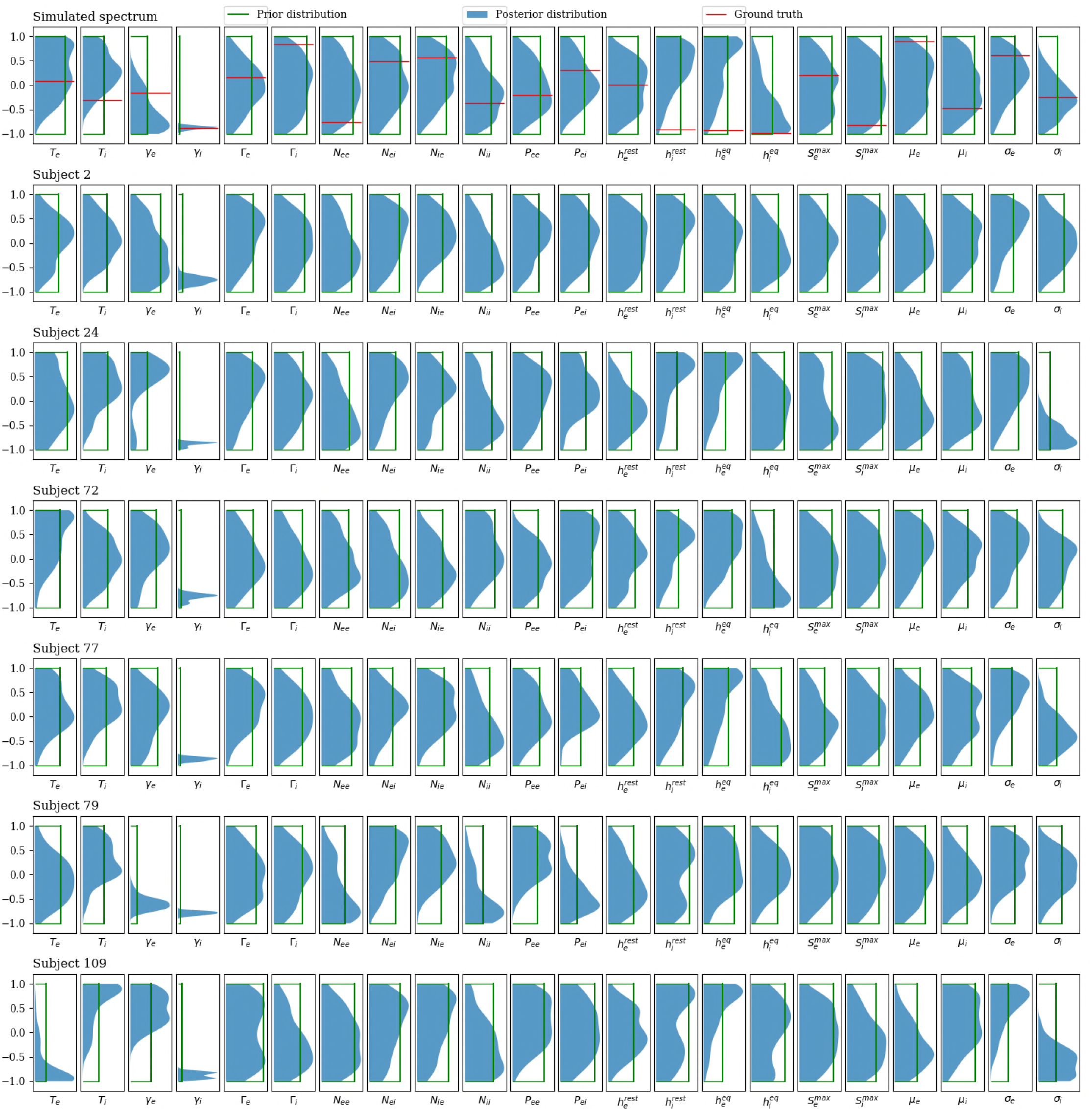
Posterior distributions based on PSO sampling. Comparison of the posterior (solid color) and prior marginal (green line) distributions for the selected subjects used in Fig 1 and Fig 2. For the simulated spectrum (first row), the distributions of the parameters are presented against the ground truths for the corresponding parameters (red line). The distributions are based on kernel density estimates from the best 100 of 1000 randomly seeded particle swarm optimizations for each subject. The seeds are uniformly distributed over the allowed parameter ranges. The major result is that, across the full set of 82 subjects, only the parameter *γ_i_* is significantly constrained. All other parameters have nearly the same uncertainties as the prior.

**Fig 4.**
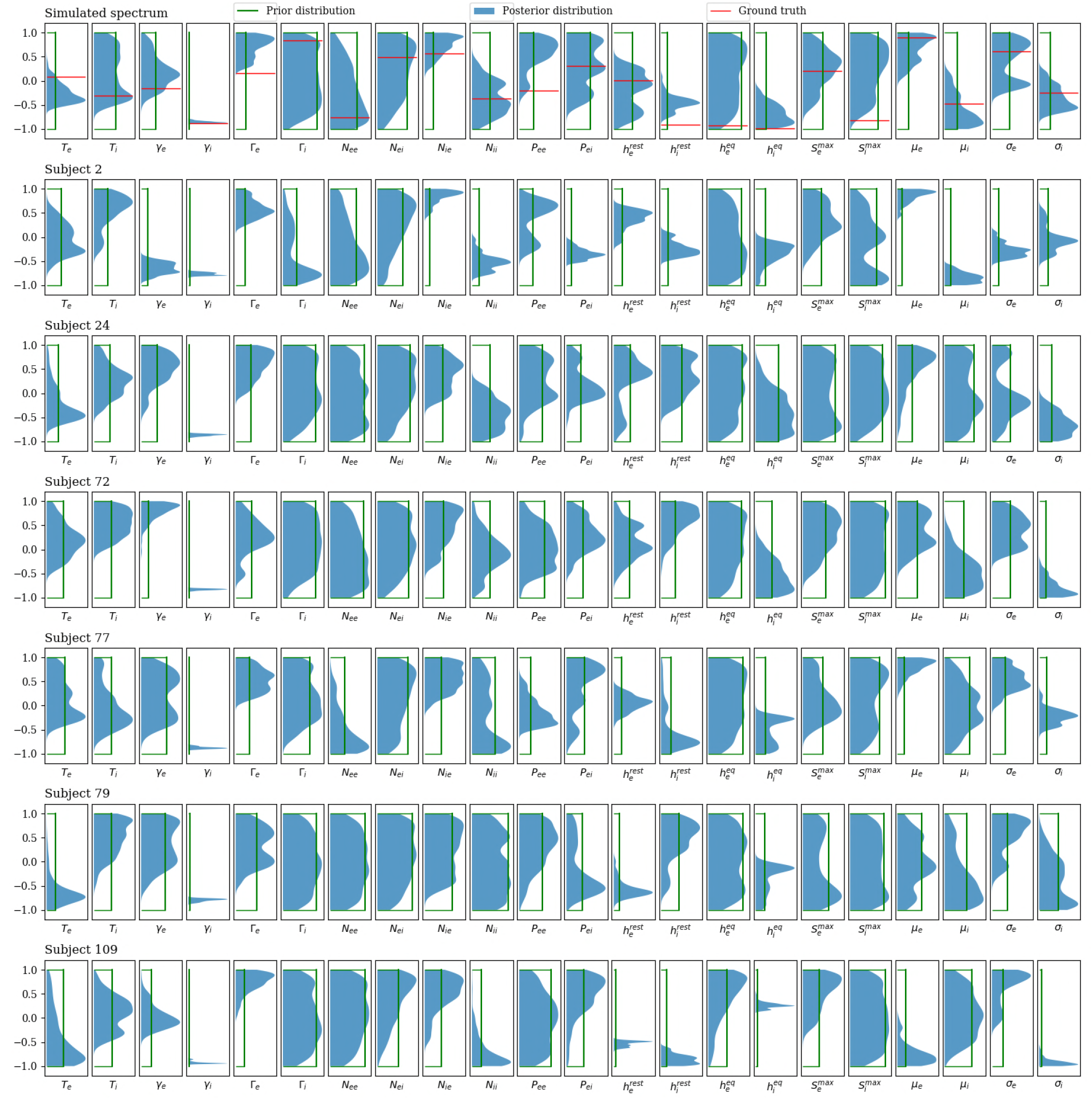
Posterior distributions based on MCMC sampling. Comparison of the posterior marginal distributions (solid color) with the prior marginal distributions (green line) for the selected subjects used in Fig 1 and Fig 2. For the simulated spectrum (first row), the distributions of the parameters are presented against the ground truths for the corresponding parameters (red line). Each distribution is based on a kernel density estimate from 1000 samples (sub-sampled from 10^6^MCMC samples). Consistent with the conclusions from PSO sampling, only *γ_i_* is consistently constrained by the data when viewed across all subjects.

**Table 1.** Physiological parameters of the model presented along with their physiological ranges. Among the 26 physiological parameters involved in the model, there are only 22 unknown parameters that are fit to the data (In our implementation, *p_ie_* and *p_ii_* are fixed constants, *γ_ee_* = *γ_ei_* ≡ *γ_e_*, and *γ_ie_* = *γ_ii_* ≡ *γ_i_*).

| No | Physiological parameter | Symbol | Min | Max |
| --- | --- | --- | --- | --- |
| 1 | Mean resting membrane potential of the excitatory population | $h_e^{rest}$ | 80 mV | 60 mV |
| 2 | Mean resting membrane potential of the inhibitory population | $h_i^{rest}$ | 80 mV | 60 mV |
| 3 | Mean Nernst membrane potential of the excitatory population | $h_e^{eq}$ | 20 mV | 10 mV |
| 4 | Mean Nernst membrane potential of the inhibitory population | $h_i^{eq}$ | 20 mV | 10 mV |
| 5 | Maximum mean firing rate of the excitatory population | $S_e^{max}$ | 0.05/ms | 0.5/ms |
| 6 | Maximum mean firing rate of the inhibitory population | $S_i^{max}$ | 0.05/ms | 0.5/ms |
| 7 | Firing thresholds of the excitatory population | $\bar{\mu}_e$ | -55 mV | -40 mV |
| 8 | Firing thresholds of the inhibitory population | $\bar{\mu}_i$ | -55 mV | -40 mV |
| 9 | Std. deviation of firing thresholds of the excitatory population | $\sigma_e$ | 2 mV | 7 mV |
| 10 | Std. deviation of firing thresholds of the inhibitory population | $\sigma_i$ | 2 mV | 7 mV |
| 11 | Passive membrane decay time const. of the excitatory population | $\tau_e$ | 5 ms | 150 ms |
| 12 | Passive membrane decay time const. of the inhibitory population | $\tau_i$ | 5 ms | 150 ms |
| 13,14 | Excitatory postsynaptic potential rate constant | $\gamma_{ee}, \gamma_{ei}$ | 0.1/ms | 1.0/ms |
| 15,16 | Inhibitory postsynaptic potential rate constant | $\gamma_{ie}, \gamma_{ii}$ | 0.01/ms | 0.5/ms |
| 17 | Postsynaptic potential amplitude of the excitatory population | $\Gamma_e$ | 0.1 mV | 0.1 mV |
| 18 | Postsynaptic potential amplitude of the inhibitory population | $\Gamma_i$ | 0.1 mV | 0.1 mV |
| 19 | Rate of the excitatory input to the excitatory population | $p_{ee}$ | 0.0 /ms | 10.0/ms |
| 20 | Rate of the excitatory input to the inhibitory population | $p_{ei}$ | 0.0 /ms | 10.0/ms |
| 21 | Rate of the inhibitory input to the excitatory population | $p_{ie}$ | 0.0/ms(fixed) | |
| 22 | Rate of the inhibitory input to the inhibitory population | $p_{ii}$ | 0.0/ms(fixed) | |
| 23 | Total number of connections an excitatory neuron receives from nearby excitatory neurons | $N_{ee}^\beta$ | 2000 | 5000 |
| 24 | Total number of connections an inhibitory neuron receives from nearby excitatory neurons | $N_{ie}^\beta$ | 100 | 1000 |
| 25 | Total number of connections an excitatory neuron receives from nearby inhibitory neurons | $N_{ei}^\beta$ | 2000 | 5000 |
| 26 | Total number of connections an inhibitory neuron receives from nearby inhibitory neurons | $N_{ii}^\beta$ | 100 | 1000 |

Among the 26 physiological parameters involved in the model, there are only 22 unknown parameters that are fit to the data (In our implementation, $p_{ie}$ and $p_{ii}$ are fixed constants, $\gamma_{ee} = \gamma_{ei} \equiv \gamma_e$ , and $\gamma_{ie} = \gamma_{ii} \equiv \gamma_i$ ).

Posterior distributions found using PSO sampling are generally broader than those found using MCMC sampling. This behavior is expected from the differences between the sampling methods: while MCMC sampling can retain correlations between samples even with significant subsampling, the different PSO samples are independent from one another. This demonstrates the superiority of the PSO approach, at least under the sampling conditions employed here. Nevertheless, both methods show that it is the postsynaptic potential rate constant of the inhibitory population, *γ_i_*, which is consistently constrained by the data across different subjects.

To better quantify how much we have learned about each parameter, the KLDs for each parameter, from all 82 subjects, are shown in Fig 5 (for PSO) and Fig 6 (for MCMC). These confirm that it is *γ_i_* that is best-constrained by the data. Most other posterior distributions are only slightly narrower than their prior distribution. Furthermore, by analysis of a simulated spectrum (see Table 2). we find that the *γ_i_* estimate is accurate as well as precise.

**Fig 5.**
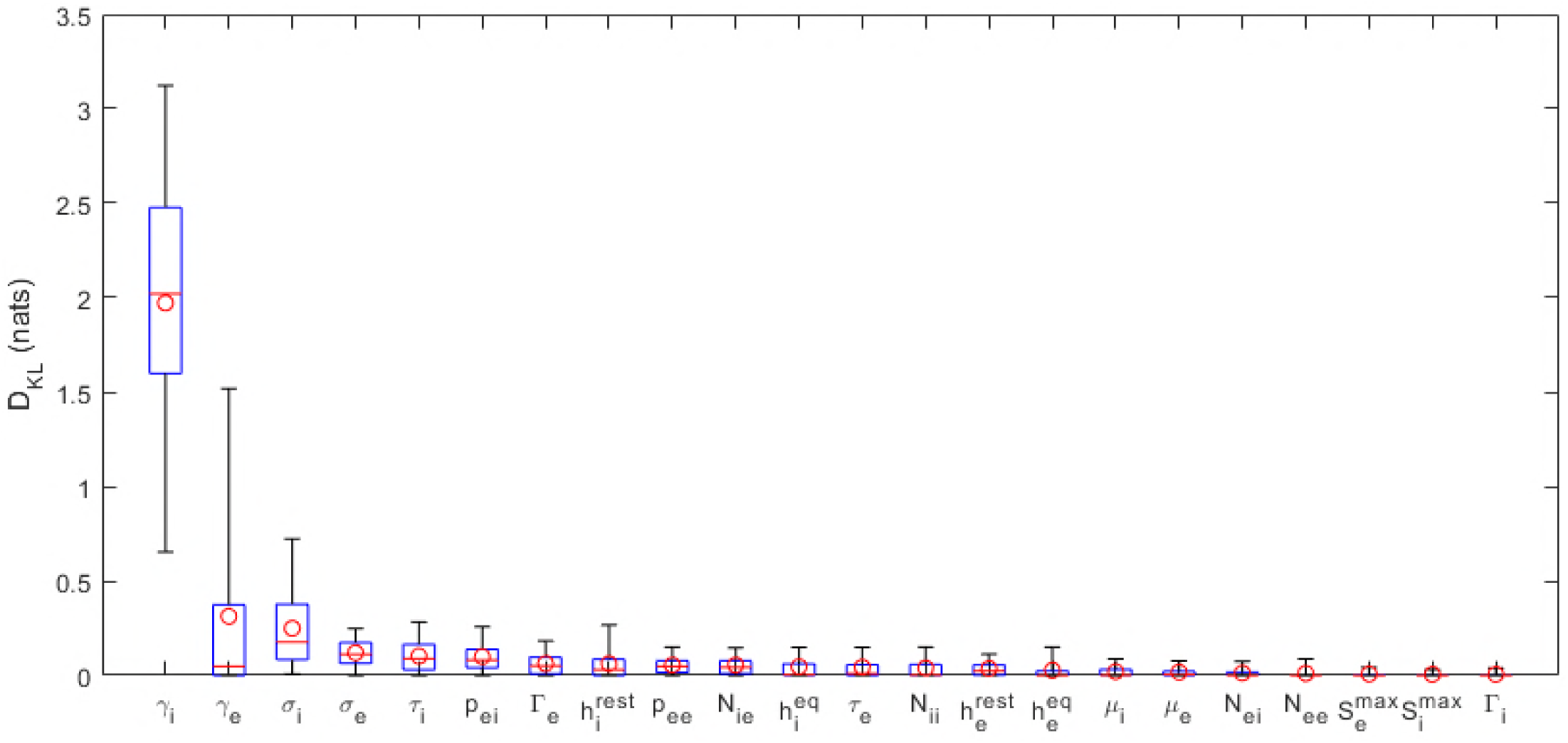
KLDs based on PSO samples. Kullback-Leibler divergences of marginal posterior parameter distributions calculated relative to uniform prior distributions. The posteriors are based on the best 100 of 1000 randomly seeded runs of particle swarm optimization (see Fig 3). The boxes represent the 25% and 75% quartiles; the whiskers represent the 5% and 95% quantiles; the red lines show the median KLDs and the circles the mean KLDs of the distributions of the KLDs over the full set of 82 subjects.

**Fig 6.**
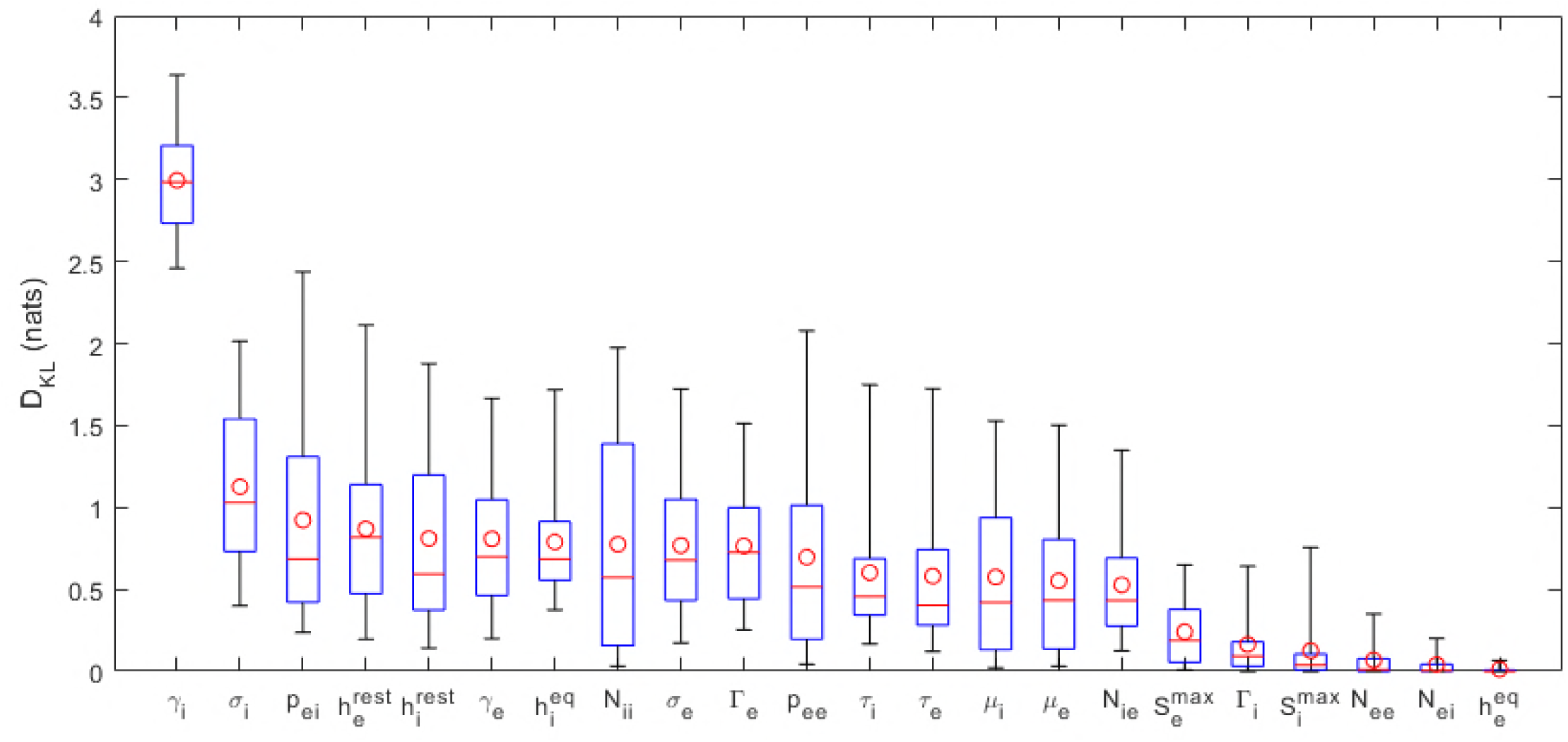
KLDs based on MCMC samples. Kullback-Leibler divergences of marginal posterior parameter distributions (see Fig 4). Here kernel density estimates based on 1000 MCMC samples of the posterior parameter distribution are used.

**Table 2.**
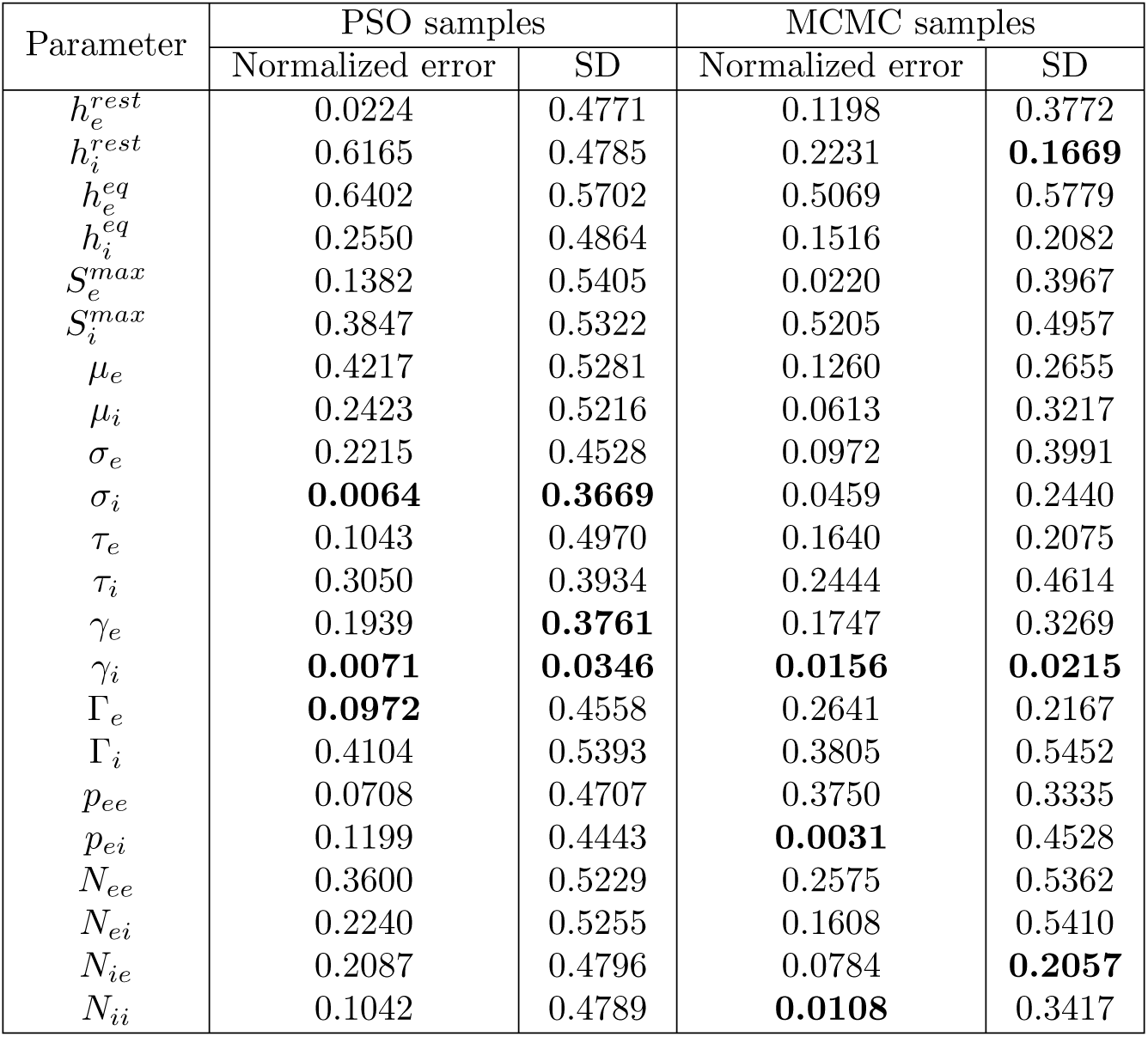
Accuracy and precision of parameter estimates in the analysis of the simulated spectrum. The accuracy of a parameter estimate is given by its deviation from the known ground truth value. Shown is the mean of this deviation for different sample estimates, normalized to the prior of the parameter. Also shown is the precision, given by the standard deviation of the sample estimates. The values in bold are the top three lowest values for each category. In this single case, the estimate of *γ_i_* always has the highest precision, for both PSO and MCMC sampling, and one of the highest accuracies.

| Parameter | PSO samples |  | MCMC samples |  |
| --- | --- | --- | --- | --- |
|  | Normalized error | SD | Normalized error | SD |
| $h_e^{rest}$ | 0.0224 | 0.4771 | 0.1198 | 0.3772 |
| $h_i^{rest}$ | 0.6165 | 0.4785 | 0.2231 | **0.1669** |
| $h_e^{eq}$ | 0.6402 | 0.5702 | 0.5069 | 0.5779 |
| $h_i^{eq}$ | 0.2550 | 0.4864 | 0.1516 | 0.2082 |
| $S_e^{max}$ | 0.1382 | 0.5405 | 0.0220 | 0.3967 |
| $S_i^{max}$ | 0.3847 | 0.5322 | 0.5205 | 0.4957 |
| $\mu_e$ | 0.4217 | 0.5281 | 0.1260 | 0.2655 |
| $\mu_i$ | 0.2423 | 0.5216 | 0.0613 | 0.3217 |
| $\sigma_e$ | 0.2215 | 0.4528 | 0.0972 | 0.3991 |
| $\sigma_i$ | **0.0064** | **0.3669** | 0.0459 | 0.2440 |
| $\tau_e$ | 0.1043 | 0.4970 | 0.1640 | 0.2075 |
| $\tau_i$ | 0.3050 | 0.3934 | 0.2444 | 0.4614 |
| $\gamma_e$ | 0.1939 | **0.3761** | 0.1747 | 0.3269 |
| $\gamma_i$ | **0.0071** | **0.0346** | **0.0156** | **0.0215** |
| $\Gamma_e$ | **0.0972** | 0.4558 | 0.2641 | 0.2167 |
| $\Gamma_i$ | 0.4104 | 0.5393 | 0.3805 | 0.5452 |
| $p_{ee}$ | 0.0708 | 0.4707 | 0.3750 | 0.3335 |
| $p_{ei}$ | 0.1199 | 0.4443 | **0.0031** | 0.4528 |
| $N_{ee}$ | 0.3600 | 0.5229 | 0.2575 | 0.5362 |
| $N_{ei}$ | 0.2240 | 0.5255 | 0.1608 | 0.5410 |
| $N_{ie}$ | 0.2087 | 0.4796 | 0.0784 | **0.2057** |
| $N_{ii}$ | 0.1042 | 0.4789 | **0.0108** | 0.3417 |

Eigenvalues of the Fisher information matrix (FIM) are shown on log scale for the selected subjects are shown in Fig 7 (for PSO) and Fig 8 (for MCMC), i.e. those computed around LS best fits and ML best fits, respectively. In all cases presented in the figures, the eigenvalues are spread over many decades with approximately uniform spacing over a log scale. This indicates the this neural population model is sloppy [36–41]. Comparison of these eigenspectra across different subjects suggests that there are usually ~5 identifiable parameter combinations for each subject.

**Fig 7.**
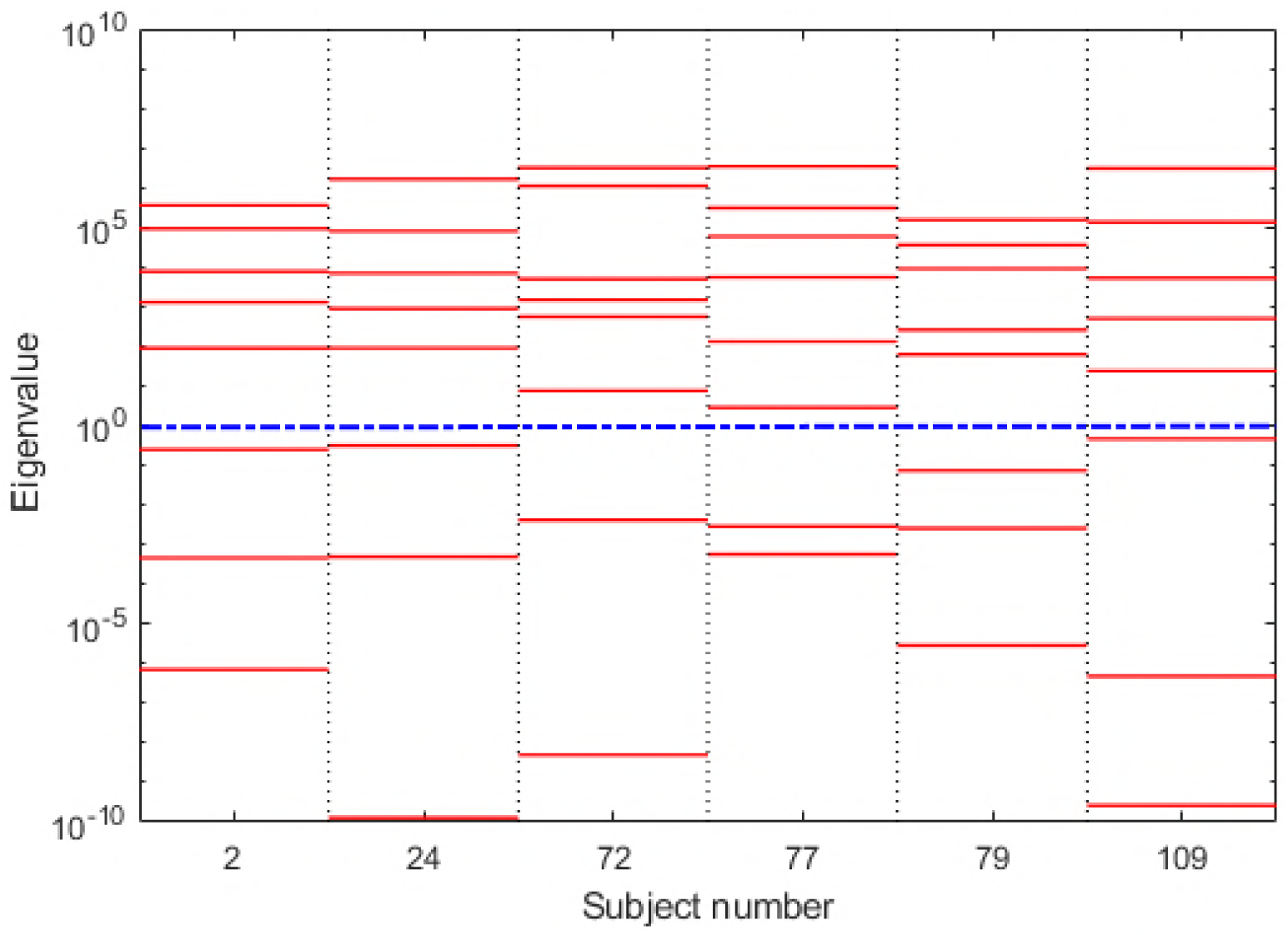
FIM eigenspectra based on LS best fits. Leading eigenvalues of the FIM for selected subjects. The FIM is numerically calculated using dimensionless increments at the parameters corresponding to a least squares fit to the experimental spectrum. Of the 22 possible eigenvalues, roughly 7 correspond to zero, at least to the numerical accuracy of the eigenvalue estimation routine. Typically 7 of the remaining 15 are too small (relative to the largest eigenvalue) to be reliably calculated using the Matlab™ eig command. The roughly uniform distribution of the eigenvalues on a log scale is a characteristic of a sloppy model. The blue dotted line delineates the separation of identifiable (above the dotted line) from unidentifiable (below the dotted line) regimes [43]. Thus ~5 parameter combinations are usually identifiable, suggesting that the 22-parameter model can be described using 5 or 6 effective parameters.

**Fig 8.**
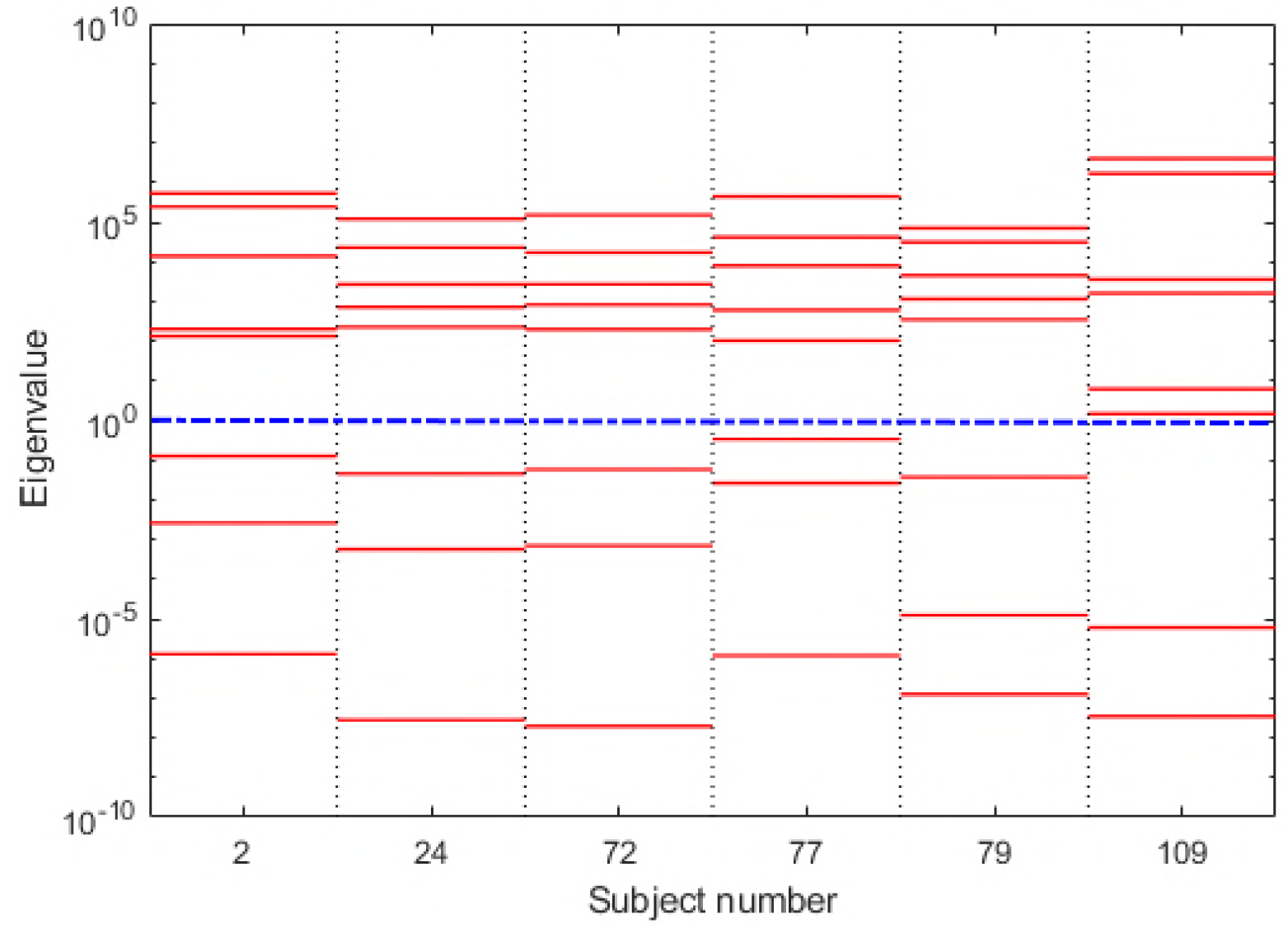
FIM eigenspectra based on ML best fits. This is similar to Fig 7, except here the FIM is calculated around the best fit found from maximum likelihood optimization for each subject’s spectrum.

The larger FIM eigenvalues define eigenvector directions corresponding to identifiable parameter combinations. To understand the parameters that contribute the most to each (identifiable) eigenvector we compute the angular distance between each parameter direction and a given eigenvector. The closer the angular distance to 0° or 180°, the greater the parameter contributes to the parameter combination and thus the more identifiable that parameter. Fig 9 (LS fits) and Fig 10 (ML fits) show the distributions, across all 82 subjects, of angular distances between each parameter and the three stiffest parameter combinations (red). For a null comparison, the angular distances to vectors randomly pointed in the 22-dimensional space are also shown (blue).

**Fig 9.**
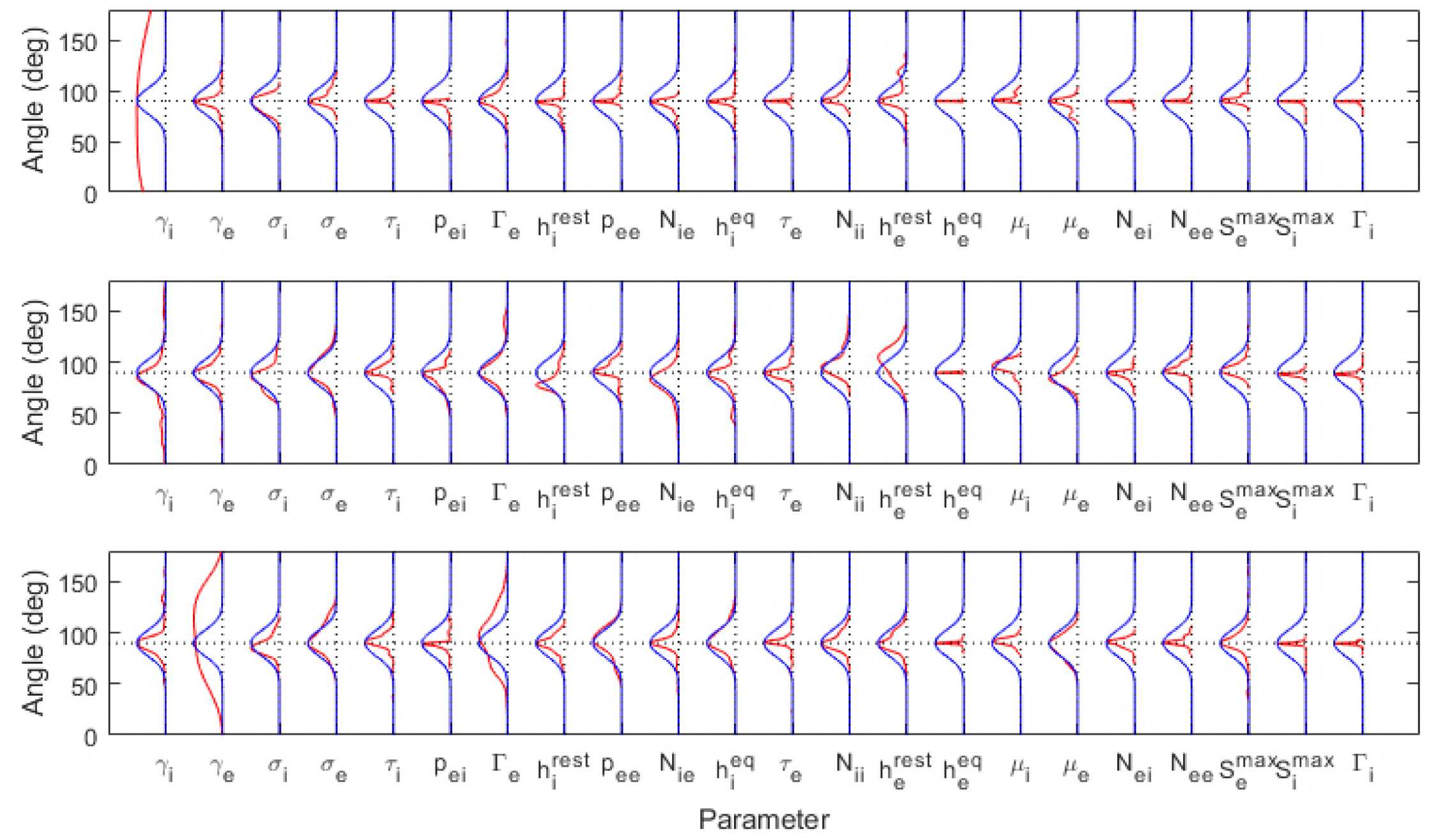
Contributions to the eigenvectors corresponding to 1*^st^*, 2*^nd^* and 3*^rd^* eigenvalues based on LS best fits. Alignment of the leading eigenvectors relative to each parameter. 0° and 180° represent perfect alignment (maximum contribution) whereas 90° represents orthogonality (no contribution). To compare the 82 subjects, results are presented as angular distributions (red lines). The first row is for the largest eigenvalue, the second row for the second-large eigenvalue, etc. The blue lines show the expected angular distributions for a randomly pointed vector in the 22-dimensional parameter space, illustrating how these are most likely to be orthogonal to any parameter direction. The angles are the inverse cosines of the direction cosines of the vectors. The distributions indicate that the parameters *γ_i_* and (to a lesser extent) *γ_e_* may play significant roles in determining the spectral form in their own right. The remaining parameters appear largely in complicated combinations.

**Fig 10.**
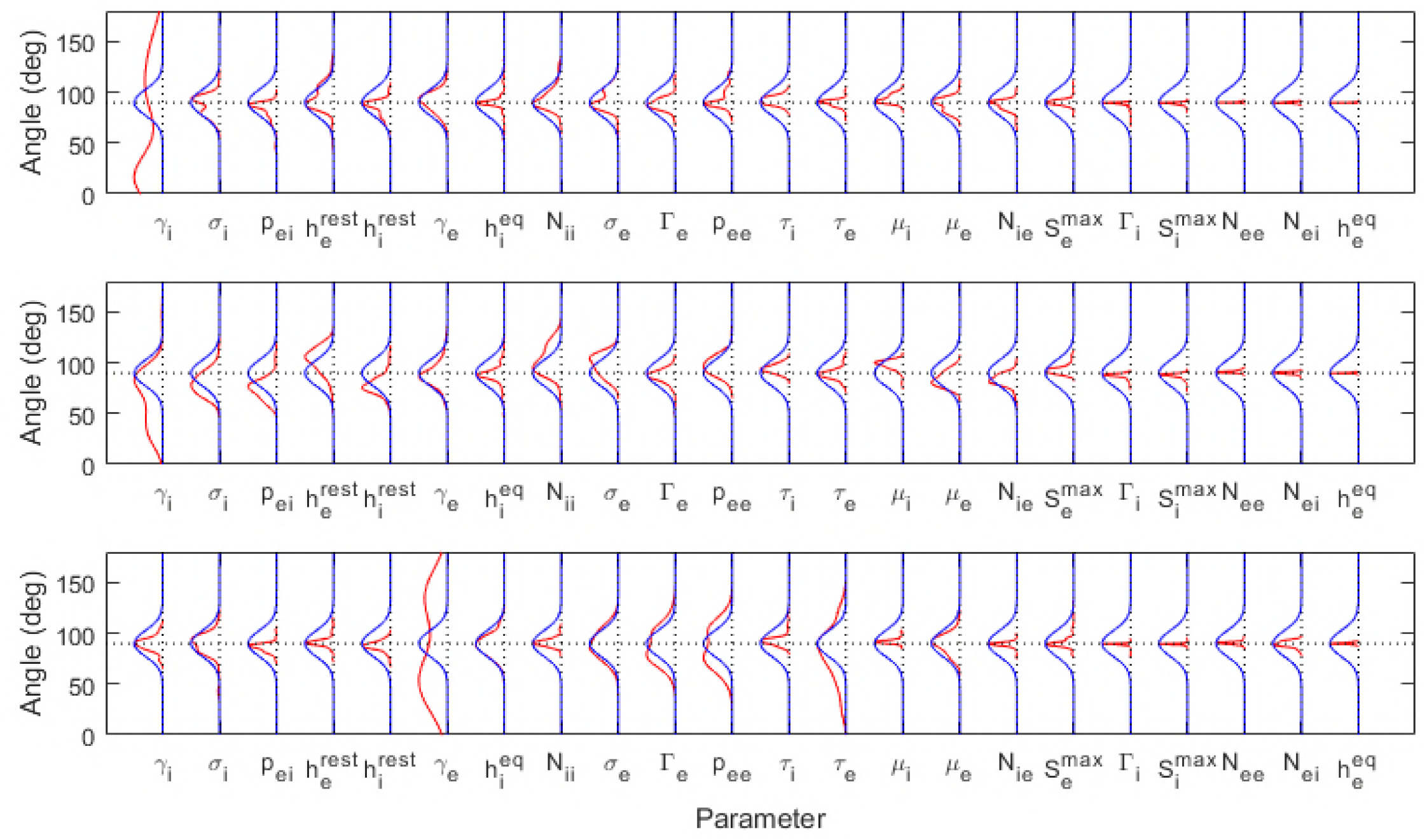
Components of eigenvectors corresponding to 1*^st^*, 2*^nd^* and 3*^rd^* eigenvalues based on ML best fits. As for Fig 9, but using the ML best fits, again showing the significant roles played by the parameters *γ_i_* and *γ_e_*.

For both LS and ML fits, *γ_i_* again stands out. It has the greatest contribution to the stiffest parameter direction, once again showing that it is identifiable. Interestingly, the postsynaptic potential rate constant of the *excitatory* population, *γ_e_*, dominates the third stiffest parameter combination. This indicates that it may also play an identifiable role in driving system dynamics, though to a lesser extent than *γ_i_*.

## Discussion

Fitting a neural population model to EEG data is an ill-posed inverse problem, where a wide range of parameter combinations are consistent with the observed spectrum (to within measurement error). Our approach to fitting such an unidentifiable model is to generate a sample of parameter estimates, all of which give a good fit to the data, and then characterize the structure of these samples. The steps we used can be summarized as follows:

1. *Optimization with a prior*: Using two different optimization methods, PSO with least squares minimization and MCMC with likelihood maximization, we search for parameter combinations which provide a good fit to the data. This required imposing a prior distribution on the parameters to ensure that only solutions within physiologically-plausible ranges were accepted.
2. *Marginal posterior: information gained about parameters individually*: To characterize the sampled parameter sets, we started by estimating the marginal posterior distribution of each parameter, using the Kullback-Leibler divergence to separate the information gained from the data from that which was already present in the prior. This showed that only *γ_i_*, the rate constant associated with inhibitory synaptic activity, had significantly less variability than its prior.
3. *Fisher information matrix: information gained about parameters collectively*: The marginal posterior characterizes our knowledge about each parameter individually. It does not capture what the data has taught us about the parameters collectively. In other words, because the data can induce correlations between parameter estimates, the marginal posteriors do not fully describe what we have learned from the data. Clearly the joint posterior distribution has the information we need, but this is difficult to estimate directly. We rely instead on the Fisher Information Matrix (FIM). The fact that many of the eigenvalues of the FIM are zero or nearly zero indicates that the manifold surrounding the global optimum is essentially flat (zero curvature) in those directions. The spacing of the first several eigenvalues is characteristic of the ‘hyper-ribbon’ geometry of sloppy models [37, 38].

Characterization of unidentifiability and sloppiness helps quantify the degree to which a model is over-parameterized. This in turn helps to illuminate how it can be simplified. The existence of correlations between parameter estimates suggests that model complexity can be reduced by grouping together, eliminating, or averaging subsets of parameters. This produces an effective model, with fewer degrees of freedom, without compromising predictive ability.

A number of model reduction techniques have been proposed for dynamical systems, such as balanced truncation [74–76], singular perturbation [77], and the manifold boundary approximation [78, 79]. In physical theory, model reduction techniques such as mean field and renormalization group methods [80] have long been used to quantify the effective parameters in complex physical systems. The concept of entropy, which enumerates the number of (unidentifiable) microstates that are consistent with a single (observable) macrostate, can be thought of as a measure of unidentifiability. Our finding that there are only ~5 identifiable eigenvalues in the FIM spectrum indicates that the number of effective parameters in our model is only about 5. The challenge is to understand what the resulting effective parameters mean physiologically.

Neural population models are coarse-grained networks composed of single neurons, where coarse-graining refers to the spatio-temporal averaging over microscopic states that occurs when performing a measurement, or calculating quantities, at macroscopic scales. When trying to interpret a coarse-grained measurement such as the EEG, our results show that even neural population models are not coarse-grained enough, since most parameters are unidentifiable. The fact that only one of the original parameters, out of 22, is consistently identifiable, a result confirmed by comparisons over 82 subjects and two different fitting routines, would seem to be a bleak result: despite the considerable effort required to fit the model, we appear to still be ignorant of 21 of the 22 parameters. However, when fitting a nonlinear model with many parameters, there is no guarantee that *any* of them will be identifiable. The fact that one has been found hints that it has a special role.

This has parallels in physical systems where the effective model parameters are the ones that remain identifiable under coarse-graining. For example, it has been shown in [39] that in diffusion processes and magnetic phase transitions, most of the microscopic parameters become unidentifiable at macroscopic scales, with only parameters such as the diffusion coefficient and average magnetization emerging unscathed. It has been suggested [81] that there may exist organizing principles that create ‘protectorates’ at mesoscopic scales, corresponding to particular parameters or parameter combinations that are robust to coarse-graining. The suggestion here then is that *γ_i_* is an effective parameter in neural population models, one that plays a central role in generating and modulating the alpha-rhythm in cortex.

We conclude by remarking that there are deep parallels between model identifiability, dynamic compensation [82–84] and evolvability [85] in a dynamical system. If the function of the system is robust, or insensitive, to changes in some of its underlying parameters, it can be impossible to infer those parameters by studying functional observables alone. Thus the study of identifiability and sloppiness is not simply a study of fitting problems but is also an examination of which parameter values are functionally essential and which are not.

## Methods of analysis

### Predicted model spectrum

The model spectrum is calculated from the spatially homogeneous version of the full model equations [11]. We make the additional assumptions that:

i. In the vicinity of the solutions corresponding to the resting eyes-open and eyes-closed spectra, the system is linearly stable.
ii. The excitatory rate constants *γ_ee_* and *γ_ei_* are equal, as are the inhibitory rate constants *γ_ii_* and *γ_ie_*.
iii. The measured EEG signal is proportional to the excitatory mean soma membrane potential, *h_e_*.
iv. The linearised system is driven by Gaussian white noise fluctuations on the external excitatory to excitatory signal *p_ee_*.

Under these assumptions, it can be shown that the linear system transfer function, *T*(*s*), is (to within an overall sign) that of a simple feedback system as shown in Fig 11 involving two third order filters:

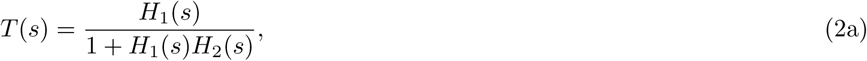

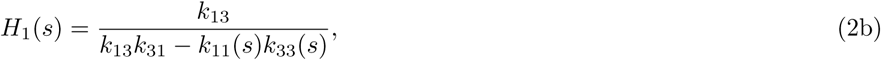

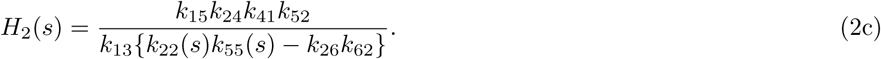

**Fig 11.**
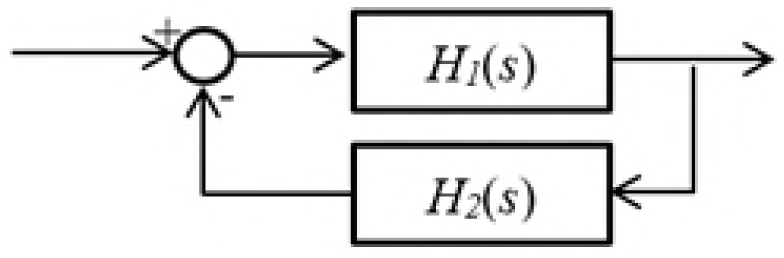
The model as a simple feedback system. The transfer function of the 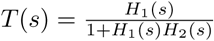where both *H*_1_(*s*) and *H*_1_(*s*) are third order filters.

The polynomials *k*_11_(*s*) and *k*_22_(*s*) are linear in *s* and *k*_33_(*s*) and *k*_55_(*s*) are quadratic in *s*. The derivation of this result and detailed expressions for the factors appearing in these equations are given in S1 Appendix.

Given that the spectra are assumed to arise from a white noise spectrum filtered by this transfer function, the expected value of the spectral estimate at frequency *ω*, given a vector of model parameters (***θ***), has the form:

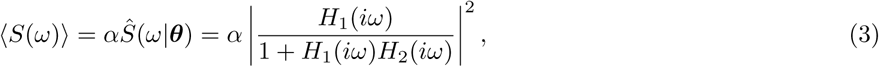

where the constant *α* accounts for the unknown driving amplitude and for attenuation due to volume conduction and other (frequency-independent) effects. The value for *α* is found using a least-squares fit to the measured spectral estimates. The analytic result is:

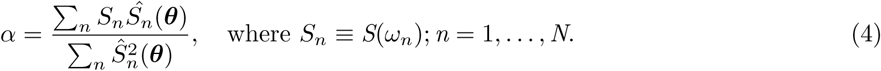

### Likelihood for the predicted model spectrum

For a spectrum of the form described above, with sufficiently high sampling rates and negligible measurement noise, the spectral estimate from the Welch periodogram at each sampled frequency {*ω_n_* = 2π*f_n_*; *n* = 1, 2, *3,…,N*} is approximately an independent random variable with a known distribution [86]. The exact form is computationally involved and for our immediate purposes we will ignore the effects of window overlap and non-uniform window shape on the resulting distributions. With this simplification the probability distribution function (pdf) for the spectral estimate, *S_n_*, is a gamma distribution:

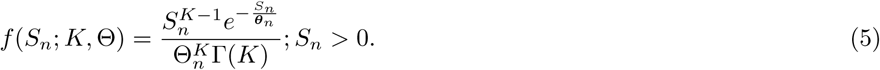

Here, for non-zero frequencies, the shape parameter *K* is found from the number of epochs averaged in the periodogram. For zero frequency, replace *K* with *K*/2 throughout. The scale parameter is given by

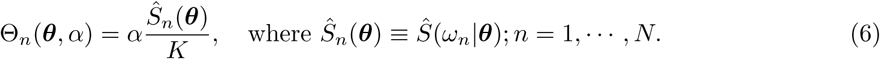

The likelihood function for the vector of spectral estimates, ***S*** = [*S*_1_ *S*_2_ *S*_3_ *… S_N_*]*^T^*, given the parameter values ***θ***, is then the product of the distributions of the individual spectral estimates:

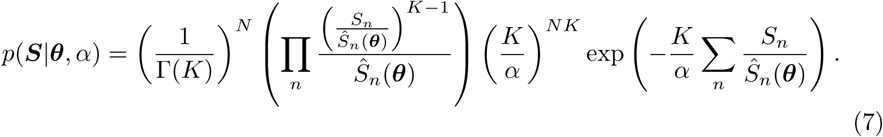

The constant *α* is adjusted to give the maximum likelihood fit of the model spectrum to the target experimental spectrum. The analytic result is that

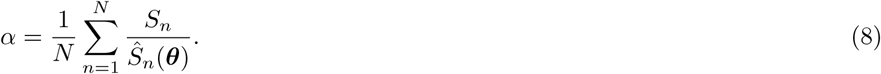

The likelihood based on model parameters alone is then

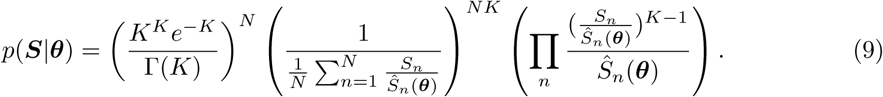

### Fitting schemes

#### Particle swarm optimization and least squares minimization

The first method we used to find the best fit parameters and their uncertainties is particle swarm optimization (PSO) [87], a standard technique for nonlinear optimization problems. PSO is an optimization algorithm inspired by swarming behavior in nature to process knowledge in the course of searching the best solution in a high-dimensional space ℝ*^D^*. At the individual level a particular particle p in a particular iteration represents a distinct candidate solution X*_p_*∈ℝ*^D^* whose quality is defined by the cost function. Throughout the iterations the particle moves around in the search-space in the direction and velocity guided by its local best known position L*_p_*∈ℝ*^D^* as well as the global best known position G∈ℝ*^D^* in such a way that in any iteration p’s velocity is given by:

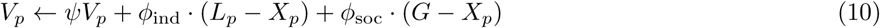

where *ψ* is a predefined inertia weight, as proposed by Shi and Eberhart [88], while 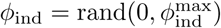and 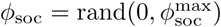are weights given to individual versus social interaction, respectively (here rand(0, *c*^max^) represents a random value between 0 and *c*^max^). The algorithm iteratively updates the global best known position G until the stopping criterion is reached.

PSO was used to estimate the 22-dimensional vector of parameters 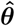that minimizes a least squares (LS) cost function *C* given by the sum of squared residuals between the measured spectrum ***S*** and the normalized model spectrum 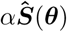:

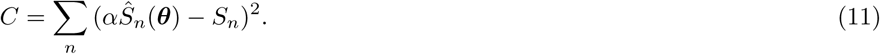

#### Markov Chain Monte Carlo and maximum likelihood estimation

In addition to the PSO method, we also used a Markov Chain Monte Carlo (MCMC) approach. For a given measured spectrum, ***S***, the best fit ***θ*** was found by maximizing the posterior distribution *p*(***θ****|****S***) given in Eq 12. To simplify the calculation, a local maximum is sought in the vicinity of the MCMC sampled parameter that maximizes the likelihood function (Eq 9) evaluated at the observed spectrum. In practice this starting value is based on the resampled parameter set rather than on the full set and the maximum found using the Matlab^®^ *fminsearch* command on the negative of the log of the posterior distribution.

### Sampling schemes

#### PSO-based random sampling

We apply an unbiased approach to draw a set of fair samples from a complicated distribution of solutions to a model-parameterization problem. Given a target spectrum to infer the distributions of the estimated parameters from, we carry out the approach by performing the three steps as follows.

1. *Generate samples*. We perform this step by using 1000 independent instantiations of the PSO to find 1000 different parameter estimates. In this scheme we use 80 particles for each swarm. Parameter starting points are chosen by random sampling from a uniform distribution over the physiologically-relevant ranges given in Table 1. During the parameter search, each parameter is forced to stay within its physiologically-relevant range by assigning a high cost to particles with values outside these ranges. This is similar to employing a Bayesian prior that is flat over the acceptable parameter range, and zero elsewhere. This step results in a preliminary set of 1000 parameter estimates 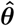.
2. *Select the best samples*. Not all samples drawn in the first step represent acceptable fits. We only accept the 10% of estimates which have the lowest cost functions, since, by inspection over fits to multiple subjects, these consistently correspond to good fits.
3. *Estimate the distributions*. Having selected samples of 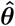, we construct kernel density estimates of the marginal posterior distributions of the parameters.

#### Markov chain Monte Carlo sampling

In the MCMC approach we employ an explicitly Bayesian framework, treating the parameters as random variables. Given a known prior distribution, 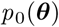, we seek the posterior distribution, conditioned on the observed spectrum ***S*** given by

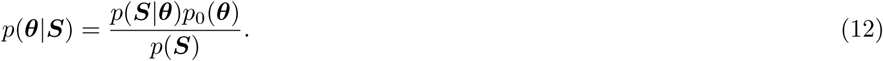

In the absence of an explicit closed form expression for this function, averaged quantities can be estimated in a Monte-Carlo fashion given a sufficiently large set of parameter values drawn randomly from this distribution. To obtain these values we use the Metropolis-Hastings Markov chain Monte Carlo (MCMC) algorithm [89] based on the log likelihood ratio that follows from the likelihood function described before.

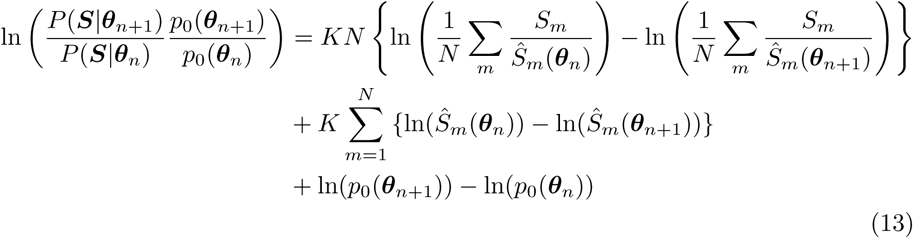

The sampled sets obtained for each target spectrum consist of 10^6^ MCMC samples with the step size (of normalised parameter values) adjusted during a burn-in phase of length 40000 to yield an acceptance ratio in the vicinity of 0.25. When appropriate, the long sampled sequence is resampled, typically to yield sequences of length 1000 upon which to base averages. For consistency with the particle swarm approach, the prior distribution is assumed to be uniform over its support.

### Kullback-Leibler divergence

A convenient measure of the information gained about individual parameters as a result of the measurement of the spectrum is the Kullback-Leibler divergence (KLD) [90, 91]. Here we use the KLD to measure the change in the marginal posterior distribution of each parameter relative to its marginal prior:

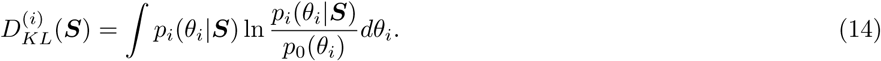

When KLDs are used to measure how posteriors differ from priors based on MCMC samples, the integral is numerically evaluated using marginal distributions approximated by kernel density estimates using 1000 parameter values resampled from the full MCMC sampled parameter set for the given spectrum. The prior distributions are uniform over their support. For consistency, the posterior kernel estimates are truncated to have the same support. The kernel density estimate for a given parameter is sampled at 100 points over its support and the integral estimated numerically. For the PSO samples, due to the limited number of independent samples, the integral is estimated using a 10 bin histogram approximation.

### Fisher information matrix

To assess the sloppiness of the model fit, we examine the eigenvalues of the Fisher information matrix (FIM), the definition of which for the pdf *P*(***S|θ***) is given by:

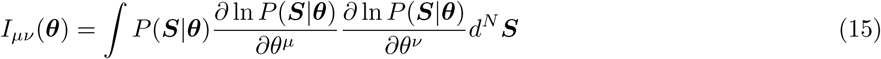

In general the integration here could present considerable difficulty, however, for the distribution given by Eq (15), it can be shown that a simplification is possible, resulting in an expression involving only the derivatives of the model spectral estimates, evaluated at the desired parameter values:

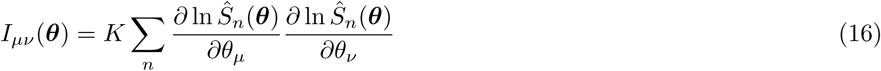

(For a derivation of this result, see S2 Appendix.)

The derivatives, with respect to normalised parameters at the LS or ML estimated values, are evaluated numerically using a 5-point finite difference approximation, and the resulting products summed over the sampled frequencies. The Matlab^®^ *eig* command is used to find the eigenvalues and eigenvectors of the resulting matrices. Numerical experiments with surrogate matrices suggest that the eigenvalues calculated using *eig* are reliable over some 10 orders of magnitude. For our modelled spectra we expect the FIM to be positive semidefinite and of less than full rank, so negative eigenvalues and eigenvalues smaller than 10^−10^ times the largest eigenvalue are taken as zero.

## Supporting information

### S1 Appendix. Derivation of the transfer function of the system

The derivation of Eq (2a) and detailed expressions for the factors appearing in Eq (2b) and Eq (2c).

### S2 Appendix. Derivation of the Fisher information matrix

The derivation of Eq (16) from Eq (15).

## Supporting information

## Acknowledgements

This work was supported in part by a Swinburne Postgraduate Research Award to AH and in part by an Australian Research Council (ARC) grant FT140101104 to DGH.

## Author contributions

**Conceptualization** DGH, PJC, DTJL; **Formal analysis** AH, PJC; **Funding acquisition** DGH; **Methodology** AH, PJC, DGH; **Software** AH, PJC; **Supervision** PJC, DTJL, DGH; **Writing - original draft preparation** AH, PJC, DTJL, DGH; **Writing - review and editing** AH, PJC, DTJL, DGH.

