## Supplementary material for "Parameter estimation and identifiability in a neural population model for electro-cortical activity"

### S1 Appendix

#### Derivation of the transfer function of the system

The full set of the Liley equations involve 14 non-linear integro-differential equations but, with reasonable assumptions about the spatial scales of the variations of the state of the system, can be reduced to an approximate set of 14 partial differential equations [Liley, Cadusch, & Wright, 1999]. For the case in which the long range inputs to the local cortical column can be treated as external given signals, typically of a stochastic nature, these can in turn be approximated by a set of 10 ordinary differential equations describing the local behaviour of the cortical column [Dafilis, Liley, & Cadusch, 2001].

In terms of symbols defined in Table 2 in the main article, these 10 equations are:

$$\tau_e \frac{dh_e}{dt} = h_{er} - h_e + \psi_{ee}(h_e) I_{ee} + \psi_{ie}(h_e) I_{ie} \tag{1a}$$

$$\tau_i \frac{dh_i}{dt} = h_{ir} - h_i + \psi_{ei}(h_i) I_{ei} + \psi_{ii}(h_i) I_{ii} \tag{1b}$$

$$\frac{dI_{ee}}{dt} = V_{ee} \tag{1c}$$

$$\frac{dV_{ee}}{dt} = -2\gamma_e V_{ee} - \gamma_e^2 I_{ee} + (\Gamma_e \gamma_e e N_{ee}^{\beta}) S_e(h_e) + (\Gamma_e \gamma_e e) p_{ee} \tag{1d}$$

$$\frac{dI_{ei}}{dt} = V_{ei} \tag{1e}$$

$$\frac{dV_{ei}}{dt} = -2\gamma_e V_{ei} - \gamma_e^2 I_{ei} + (\Gamma_e \gamma_e e N_{ei}^{\beta}) S_e(h_e) + (\Gamma_e \gamma_e e) p_{ei} \tag{1f}$$

$$\frac{dI_{ie}}{dt} = V_{ie} \tag{1g}$$

$$\frac{dV_{ie}}{dt} = -2\gamma_i V_{ie} - \gamma_i^2 I_{ie} + (\Gamma_i \gamma_i e N_{ie}^{\beta}) S_i(h_i) + (\Gamma_i \gamma_i e) p_{ie} \tag{1h}$$

$$\frac{dI_{ii}}{dt} = V_{ii} \tag{1i}$$

$$\frac{dV_{ii}}{dt} = -2\gamma_i V_{ii} - \gamma_i^2 I_{ii} + (\Gamma_i \gamma_i e N_{ii}^{\beta}) S_i(h_i) + (\Gamma_i \gamma_i e) p_{ii} \tag{1j}$$

where

$$\psi_{ee}(h_e) = \frac{h_e^{eq} - h_e}{|h_e^{eq} - h_e^{rest}|} \tag{1k}$$

$$\psi_{ie}(h_e) = \frac{h_i^{eq} - h_e}{|h_i^{eq} - h_e^{rest}|} \tag{1l}$$

$$\psi_{ei}(h_i) = \frac{h_e^{eq} - h_i}{|h_e^{eq} - h_i^{rest}|} \tag{1m}$$

$$\psi_{ii}(h_i) = \frac{h_i^{eq} - h_i}{|h_i^{eq} - h_i^{rest}|} \tag{1n}$$

To linearize these we restrict attention to fixed points that correspond to stable point attractors. The fixed point equations can be reduced to:

$$V_{ee} = V_{ei} = V_{ie} = V_{ii} = 0 \tag{2a}$$

$$\bar{I}_{ee} = \frac{\Gamma_e e}{\gamma_e}[N_{ee}^{\beta}]S_e(\bar{h}_e) + \left(\frac{\Gamma_e e}{\gamma_e}\right)\bar{p}_{ee} \tag{2b}$$

$$\bar{I}_{ei} = \frac{\Gamma_e e}{\gamma_e}[N_{ei}^{\beta}]S_e(\bar{h}_e) + \left(\frac{\Gamma_e e}{\gamma_e}\right)\bar{p}_{ei} \tag{2c}$$

$$\bar{I}_{ie} = \left(\frac{\Gamma_i e N_{ie}^{\beta}}{\gamma_i}\right)S_i(\bar{h}_i) + \left(\frac{\Gamma_i e}{\gamma_i}\right)\bar{p}_{ie} \tag{2d}$$

$$\bar{I}_{ii} = \left(\frac{\Gamma_i e N_{ii}^{\beta}}{\gamma_i}\right)S_i(\bar{h}_i) + \left(\frac{\Gamma_i e}{\gamma_i}\right)\bar{p}_{ii} \tag{2e}$$

$$0 = h_{er} - \bar{h}_e + \psi_{ee}(\bar{h}_e)\bar{I}_{ee} + \psi_{ie}(\bar{h}_e)\bar{I}_{ie} \tag{2f}$$

$$0 = h_{ir} - \bar{h}_i + \psi_{ei}(\bar{h}_i)\bar{I}_{ei} + \psi_{ii}(\bar{h}_i)\bar{I}_{ii} \tag{2g}$$

where the overbar on a state variable denotes its fixed point value and on an input variable ($p_{ee}, p_{ei}$ etc.) denotes its DC value, treated as a tonic system parameter.

The linearized equations are obtained by expanding the state variables around their fixed point values; the non-zero terms of the corresponding Jacobian, evaluated at the fixed point, are of the form:

$$J_{11} = \frac{1}{\tau_e}\left(-1 - \frac{\bar{I}_{ee}}{|(h_e^{eq} - h_e^{rest}|} - \frac{\bar{I}_{ie}}{|h_i^{eq} - h_e^{rest}|}\right) \tag{3a}$$
$$J_{13} = \frac{\psi_{ee}(\bar{h}_e)}{\tau_e} \tag{3b}$$
$$J_{17} = \frac{\psi_{ie}(\bar{h}_e)}{\tau_e} \tag{3c}$$
$$J_{22} = \frac{1}{\tau_i}\left(-1 - \frac{\bar{I}_{ei}}{|h_e^{eq} - h_i^{rest}|} - \frac{\bar{I}_{ii}}{|h_i^{eq} - h_i^{rest}|}\right) \tag{3d}$$
$$J_{25} = \frac{\psi_{ei}(\bar{h}_i)}{\tau_i} \tag{3e}$$
$$J_{29} = \frac{\psi_{ii}(\bar{h}_i)}{\tau_i} \tag{3f}$$
$$J_{41} = \Gamma_e \gamma_e e N_{ee}^{\beta} S_e^{'}(\bar{h}_e) \tag{3g}$$
$$J_{43} = -\gamma_e^2 \tag{3h}$$
$$J_{44} = -2\gamma_e \tag{3i}$$
$$J_{61} = \Gamma_e \gamma_e e N_{ei}^{\beta} S_e^{'}(\bar{h}_e) \tag{3j}$$
$$J_{65} = -\gamma_e^2 \tag{3k}$$
$$J_{66} = -2\gamma_e \tag{3l}$$
$$J_{82} = \Gamma_i \gamma_i e N_{ie}^{\beta} S_i^{'}(\bar{h}_i) \tag{3m}$$
$$J_{87} = -\gamma_i^2 \tag{3n}$$
$$J_{10,2} = \Gamma_i \gamma_i e N_{ii}^{\beta} S_i^{'}(\bar{h}_i) \tag{3o}$$
$$J_{10,9} = -\gamma_i^2 \tag{3p}$$
$$J_{10,10} = -2\gamma_i \tag{3q}$$
$$J_{34} = J_{56} = J_{78} = J_{9,10} = 1 \tag{3r}$$

The symbol $S_e^{'}(\bar{h}_e)$ denotes the derivative of $S_e(\bar{h}_e)$ w.r.t. $h_e$ evaluated at the fixed point value $\bar{h}_e$, and similarly for $S_i^{'}(\bar{h}_i)$. Since $J_{43} = J_{65}$ and $J_{87} = J_{10,9}$ and $J_{43} = -\left(\frac{J_{44}}{2}\right)^2$ and $J_{87} = -\left(\frac{J_{88}}{2}\right)^2$ there are only 12 independent elements of the Jacobian viz: the (1,1), (1,3), (1,7), (2,2), (2,5), (2,9), (4,1), (4,4), (6,1), (8,2), (8,8) and (10,2) elements. The resulting structure of the Jacobian is:

$$\begin{bmatrix} J_{11} & 0 & J_{13} & 0 & 0 & 0 & J_{17} & 0 & 0 & 0 \\ 0 & J_{22} & 0 & 0 & J_{25} & 0 & 0 & 0 & J_{29} & 0 \\ 0 & 0 & 0 & 1 & 0 & 0 & 0 & 0 & 0 & 0 \\ J_{41} & 0 & J_{43} & J_{44} & 0 & 0 & 0 & 0 & 0 & 0 \\ 0 & 0 & 0 & 0 & 1 & 0 & 0 & 0 & 0 & 0 \\ J_{61} & 0 & 0 & 0 & J_{65} & J_{66} & 0 & 0 & 0 & 0 \\ 0 & 0 & 0 & 0 & 0 & 0 & 0 & 1 & 0 & 0 \\ 0 & J_{82} & 0 & 0 & 0 & 0 & J_{87} & J_{88} & 0 & 0 \\ 0 & 0 & 0 & 0 & 0 & 0 & 0 & 0 & 0 & 1 \\ 0 & J_{10,2} & 0 & 0 & 0 & 0 & 0 & 0 & J_{10,9} & J_{10,10} \end{bmatrix} \tag{4}$$

The system transfer function can be obtained from the Jacobian in this form, but it is more convenient to rewrite the 10 first-order equations as a set of 2 first order equations and 4 second order equations. It is also convenient to rewrite them in terms of lumped parameters. The resulting equations are:

$$\tau_e \frac{dh_e}{dt} = h_{er} - h_e + \psi_{ee}(h_e)I_{ee} + \psi_{ie}(h_e)I_{ie} \tag{5a}$$

$$\tau_i \frac{dh_i}{dt} = h_{ir} - h_i + \psi_{ei}(h_i)I_{ei} + \psi_{ii}(h_i)I_{ii} \tag{5b}$$

$$\left(\frac{d}{dt} + \gamma_e\right)^2 I_{ee} = A_{ee}S_e(h_e) + B_{ee} + u_{ee}(t) \tag{5c}$$

$$\left(\frac{d}{dt} + \gamma_e\right)^2 I_{ei} = A_{ei}S_e(h_e) + B_{ei} + u_{ei}(t) \tag{5d}$$

$$\left(\frac{d}{dt} + \gamma_i\right)^2 I_{ie} = A_{ie}S_i(h_i) + B_{ie} + u_{ie}(t) \tag{5e}$$

$$\left(\frac{d}{dt} + \gamma_i\right)^2 I_{ii} = A_{ii}S_i(h_i) + B_{ii} + u_{ii}(t) \tag{5f}$$

The lumped parameters are

$$A_{ee} = (\Gamma_e \gamma_e e N_{ee}^{\beta}) \tag{6a}$$

$$B_{ee} = (\Gamma_e \gamma_e e)\bar{p}_{ee} \tag{6b}$$

$$A_{ei} = (\Gamma_e \gamma_e e N_{ei}^{\beta}) \tag{6c}$$

$$B_{ei} = (\Gamma_e \gamma_e e)\bar{p}_{ei} \tag{6d}$$

$$A_{ie} = (\Gamma_i \gamma_i e N_{ie}^{\beta}) \tag{6e}$$

$$B_{ie} = (\Gamma_i \gamma_i e)\bar{p}_{ie} \tag{6f}$$

$$A_{ii} = (\Gamma_i \gamma_i e N_{ii}^{\beta}) \tag{6g}$$

$$B_{ii} = (\Gamma_i \gamma_i e)\bar{p}_{ii} \tag{6h}$$

The $u_{ee}(t)$ etc. are the time varying parts of the inputs to the system. With our conventional assumptions the linearized equations involve 12 distinct parameters or combinations of system parameters, viz: $\tau_e$, $\tau_i$,$\gamma_e$, $\gamma_i$, and the quantities $A_{ab}S_a^{'}(\bar{h}_a)$ and $B_{ab}$. In practice, however, we usually assume (on physiological grounds) that $B_{ii} = B_{ie} = 0$, reducing the lumped parameter set to 10 distinct elements compared to the 22 independent "physiological" parameters of the 10D system. Typically the dominant input to a cortical column is ecitatory to excitatory, so in the following we also assume that $u_{ei}(t) = u_{ie}(t) = u_{ii}(t) = 0$. Also on physiological grounds, the output of the system (the local EEG signal) is assumed to be proportional to $h_e(t) - \bar{h}_e$.

#### The 10D system spectrum.

The 10D equations with the coefficient assignments above lead to a simplification of the spectrum. Using the mixed first and second order form, the Laplace transformed linearized equations become:

$$K(s)\mathbf{y} = \mathbf{u} \tag{7}$$

where $\mathbf{y}$ is the vector of the deviations of the state variables from their fixed point values and $\mathbf{u}$ is the vector of time varying parts of the input signals and K is a $6 \times 6$

matrix:

$$K(s) = \begin{bmatrix} k_{11}(s) & 0 & k_{13} & 0 & k_{15} & 0 \\ 0 & k_{22}(s) & 0 & k_{24} & 0 & k_{26} \\ k_{31} & 0 & k_{33}(s) & 0 & 0 & 0 \\ k_{41} & 0 & 0 & k_{33}(s) & 0 & 0 \\ 0 & k_{52} & 0 & 0 & k_{55}(s) & 0 \\ 0 & k_{62} & 0 & 0 & 0 & k_{55}(s) \end{bmatrix} \tag{8}$$

with

$$k_{11}(s) = s + \frac{1}{\tau_e}\left(1 + \frac{\bar{I}_{ee}}{|h_e^{eq} - h_e^{rest}|} + \frac{\bar{I}_{ie}}{|h_i^{eq} - h_e^{rest}|}\right) \tag{9a}$$

$$k_{13} = -\frac{\psi_{ee}(\bar{h}_e)}{\tau_e} \tag{9b}$$

$$k_{15} = -\frac{\psi_{ie}(\bar{h}_e)}{\tau_e} \tag{9c}$$

$$k_{22}(s) = s + \frac{1}{\tau_i}\left(1 + \frac{\bar{I}_{ei}}{|h_e^{eq} - h_i^{rest}|} + \frac{\bar{I}_{ii}}{|h_i^{eq} - h_i^{rest}|}\right) \tag{9d}$$

$$k_{24} = -\frac{\psi_{ei}(\bar{h}_i)}{\tau_i} \tag{9e}$$

$$k_{26} = -\frac{\psi_{ii}(\bar{h}_i)}{\tau_i} \tag{9f}$$

$$k_{31} = -A_{ee} S_e^{'}(\bar{h}_e) \tag{9g}$$

$$k_{33}(s) = (s + \gamma_e)^2 \tag{9h}$$

$$k_{41} = -A_{ei} S_e^{'}(\bar{h}_e) \tag{9i}$$

$$k_{52} = -A_{ie} S_i^{'}(\bar{h}_i) \tag{9j}$$

$$k_{55}(s) = (s + \gamma_i)^2 \tag{9k}$$

$$k_{62} = -A_{ii} S_i^{'}(\bar{h}_i) \tag{9l}$$

The spectrum of interest is proportional to the (1,3) element of the inverse of $K$ evaluated along the imaginary axis of $s$. This can be written as the quotient:

$$\hat{S}(\omega) = \left|\frac{\det(H(i\omega)}{\det(K(i\omega)}\right|^2 \tag{10}$$

where $H$ is the (3,1) minor of $K$:

$$H(s) = \begin{bmatrix} 0 & k_{13} & 0 & k_{15} & 0 \\ k_{22}(s) & 0 & k_{24} & 0 & k_{26} \\ 0 & 0 & k_{33}(s) & 0 & 0 \\ k_{52} & 0 & 0 & k_{55} & 0 \\ k_{62} & 0 & 0 & 0 & k_{55}(s) \end{bmatrix} \tag{11}$$

Now, using Matlab for symbolic manipulations, we get:

$$\det K = -k_{33}(s)k_{55}(s)[\{k_{22}(s)k_{55}(s) - k_{26}k_{62}\}\{k_{13}k_{31} - k_{11}(s)k_{33}(s)\} + k_{15}k_{24}k_{41}k_{52}]$$

$$ \tag{12} $$

and

$$\det H = -k_{13}k_{33}(s)k_{55}(s)\{k_{22}(s)k_{55}(s) - k_{26}k_{62}\} \tag{13}$$

In $\hat{S}(\omega)$, the second order zeros $k_{33}(s)$ and $k_{55}(s)$ in det $H$ cancel the corresponding second order poles in det $K$, leading to

$$\hat{S}(\omega) = \left| \frac{k_{13}\{k_{22}(i\omega)k_{55}(i\omega) - k_{26}k_{62}\}}{[\{k_{22}(i\omega)k_{55}(i\omega) - k_{26}k_{62}\}\{k_{13}k_{31} - k_{11}(i\omega)k_{33}(i\omega)\} + k_{15}k_{24}k_{41}k_{52}]} \right|^2 \tag{14}$$

So, consistent with the numerical results, the spectrum is a rational form with 6 poles and 3 zeros rather than 10 poles and 7 zeros as might be expected from simple power counting.

The structure of the transfer function, $T(s)$, is (to within an overall sign) that of a simple feedback system involving two third order filters:

$$T(s) = \frac{H_1(s)}{1 + H_1(s)H_2(s)} \tag{15a}$$

$$H_1(s) = \frac{k_{13}}{k_{13}k_{31} - k_{11}(s)k_{33}(s)} \tag{15b}$$

$$H_2(s) = \frac{k_{15}k_{24}k_{41}k_{52}}{k_{13}\{k_{22}(s)k_{55}(s) - k_{26}k_{62}\}} \tag{15c}$$

$$\hat{S}(\omega) = |T(i\omega)|^2 \tag{15d}$$

In this case, the pole locations of $H_1$ are governed mainly by properties of excitatory cells while those of $H_2$ by the properties of inhibitory cells. In practice, $H_1$ appears to be a low-pass filter and $H_2$ is resonant.

### S2 Appendix

#### Derivation of the Fisher Information Matrix

The definition of the Fisher information matrix (FIM) for a p.d.f. $f(\mathbf{x}|\boldsymbol{\theta})$ is :

$$I_{\mu\nu}(\boldsymbol{\theta}) = \int f(\mathbf{x}|\boldsymbol{\theta}) \frac{\partial \ln f(\mathbf{x}|\boldsymbol{\theta})}{\partial \theta_\mu} \frac{\partial \ln f(\mathbf{x}|\boldsymbol{\theta})}{\partial \theta_\nu} d^N\mathbf{x} \tag{1}$$

If the components of the vector $\mathbf{x}$ are independent then

$$f(\mathbf{x}|\boldsymbol{\theta}) = \prod_k f_k(x_k|\boldsymbol{\theta}) \tag{2}$$

For this case, taking account of the normalisation, $\int f_k(x_k|\boldsymbol{\theta})dx_k = 1$, it is straightforward to show that:

$$I_{\mu\nu}(\boldsymbol{\theta}) = \sum_k I^{(k)}_{\mu\nu}(\boldsymbol{\theta}) \tag{3}$$

where

$$I^{(k)}_{\mu\nu}(\boldsymbol{\theta}) = \int f_k(x_k|\boldsymbol{\theta}) \frac{\partial \ln f_k(x_k|\boldsymbol{\theta})}{\partial \theta_\mu} \frac{\partial \ln f_k(x_k|\boldsymbol{\theta})}{\partial \theta_\nu} dx_k \tag{4}$$

is the FIM based on the marginal p.d.f. for the component $x_k$.

For the problem in which the components of the vector $\mathbf{x}$ are model spectral estimates at different frequencies evaluated using the Welch periodogram, and providing that the correlations introduced by the overlap and non-uniform shapes of the windows used can be neglected, the spectral estimate at a given frequency has a gamma distribution of the form:

$$f_k(x_k; K, \Theta_k) = \frac{x_k^{K-1} e^{-\frac{x_k}{\Theta_k}}}{\Theta_k^K \Gamma(K)}; x_k \geq 0 \tag{5}$$

where $K$ is the number of segments averaged in the Welch periodogram (which is independent of the system parameters, $\boldsymbol{\theta}$) and the scale parameter, $\Theta_k$, (which does depend on the system parameters) is related to the model's expected power spectral density at the given frequency by

$$\Theta_k = \frac{\alpha |T(i\omega_k|\boldsymbol{\theta})|^2}{K} \tag{6}$$

In this equation, $T(s|\boldsymbol{\theta})$ is the linearized model transfer function, $\omega_k$ is the $k^{th}$ sampled frequency and $\alpha$ is the power matching parameter introduced in the main article. For our immediate purpose, investigating the sensitivity of the model output to changes in the system parameters, we can treat $\alpha$ as a given constant. With these distributions the integrals in the definition of the Fisher information matrix can be explicitly performed and the expression for $I^{(k)}_{\mu\nu}(\boldsymbol{\theta})$ considerably simplified. Thus (dropping the $k$ subscripts for convenience):

$$\ln f(x; K, \Theta) = (K-1)\ln x - \frac{x}{\Theta} - K \ln \Theta - \ln \Gamma(K) \tag{7}$$

so

$$\frac{\partial}{\partial \theta_\mu} \ln f(x; K, \Theta) = (x - K\Theta) \frac{1}{\Theta^2} \frac{\partial \Theta}{\partial \theta_\mu} \tag{8}$$

Then, substituting for the derivatives and moving terms independent of $x$ outside the integral, we get

$$I_{\mu\nu}(\boldsymbol{\theta}) = \frac{1}{\Theta^4}\frac{\partial\Theta}{\partial\theta_\mu}\frac{\partial\Theta}{\partial\theta_\nu}\int f(x;K,\Theta)(x-K\Theta)^2 dx \tag{9}$$

The mean of the gamma distribution is $K\Theta$ so the integral in the equation is equal to the variance of the gamma distribution, which is equal to $K\Theta^2$, thus:

$$\begin{aligned} I_{\mu\nu}(\boldsymbol{\theta}) &= \frac{1}{\Theta^4}\frac{\partial\Theta}{\partial\theta_\mu}\frac{\partial\Theta}{\partial\theta_\nu}\ \mathrm{var}(X) \\ &= \frac{K}{\Theta^2}\frac{\partial\Theta}{\partial\theta_\mu}\frac{\partial\Theta}{\partial\theta_\nu} \\ &= K\frac{\partial\ln\Theta}{\partial\theta_\mu}\frac{\partial\ln\Theta}{\partial\theta_\nu} \\ &= K\frac{\partial\ln|T(i\omega|\boldsymbol{\theta})|^2}{\partial\theta_\mu}\frac{\partial\ln|T(i\omega|\boldsymbol{\theta})|^2}{\partial\theta_\nu} \end{aligned} \tag{10}$$

Thus, restoring the $k$ subscripts and defining the sampled values of the model power spectral density, $\hat{S}_k \equiv |T(i\omega_k|\boldsymbol{\theta})|^2$, we get

$$I_{\mu\nu}(\boldsymbol{\theta}) = K\sum_k \frac{\partial\ln\hat{S}_k}{\partial\theta_\mu}\frac{\partial\ln\hat{S}_k}{\partial\theta_\nu} \tag{11}$$

Since, in the region of interest, the transfer function for the linearized model appears to have no zeros on the imaginary axis, it follows that $\hat{S}_k > 0$ and this form provides a useful simplification of the FIM.
